# Multilocus adaptation to vaccination

**DOI:** 10.1101/2021.06.01.446592

**Authors:** David V. McLeod, Sylvain Gandon

## Abstract

Pathogen adaptation to public health interventions, such as vaccination, may take tortuous routes and involve multiple mutations at distinct locations in the pathogen genome, acting on distinct phenotypic traits. Despite its importance for public health, how these multilocus adaptations jointly evolve is poorly understood. Here we consider the joint evolution of two adaptations: the pathogen’s ability to escape the vaccine-induced immune response and adjustments to the pathogen’s virulence and transmissibility. We elucidate the role played by epistasis and recombination, with an emphasis on the different protective effects of vaccination. We show that vaccines reducing transmission and/or increasing clearance generate positive epistasis between the vaccine-escape and virulence alleles, favouring strains that carry both mutations, whereas vaccines reducing virulence mortality generate negative epistasis, favouring strains that carry either mutation, but not both. High rates of recombination can affect these predictions. If epistasis is positive, frequent recombination can lead to the sequential fixation of the two mutations and prevent the transient build-up of more virulent escape strains. If epistasis is negative, frequent recombination between loci can create an evolutionary bistability, such that whichever adaptation is more accessible tends to be favoured in the long-term. Our work provides a timely alternative to the variant-centered perspective on pathogen adaptation and captures the effect of different types of vaccines on the interference between multiple adaptive mutations.

## Introduction

Pathogen evolution in response to public health interventions, such as vaccines or antimicrobials, is often complex, involving multiple genetic mutations affecting distinct aspects of pathogen life-history. For instance, a mutation at one locus may facilitate the escape of vaccine protections, or render an antibiotic ineffective, but carry a fitness cost, while a second, compensatory mutation at a different locus may reduce this cost. Crucially, any such fitness interaction between mutations (i.e., epistasis) means that the transient evolutionary dynamics of each individual mutation will hinge upon their non-random associations with the other (i.e., linkage disequilibrium), and so cannot be easily disentangled. Such multilocus dynamics are especially relevant for vaccination: although vaccines are a comparatively evolutionarily-robust public health measure [1–3], pathogen evolution to vaccination can occur, with two key adaptive routes identified.

First, pathogens can undergo rapid antigenic evolution, allowing them to escape the vaccine-induced immune response (vaccine-escape). This not only presents a barrier to developing effective vaccines [4], but it may require that vaccines be regularly updated to match with circulating antigens, a time consuming process [5], with substantial forecasting uncertainty [6, 7]. Rapid antigenic evolution has led to a focus upon developing ‘universal’ vaccines, which target conserved viral sequences and so are expected to provide more robust protection [8–10]. Second, by altering the selective balance between transmission and virulence, vaccination can select for more virulent pathogens [11–16]. Specifically, if vaccines are imperfect and so do not completely block infection, the pathogen population experiences two selective environments, vaccinated and unvaccinated hosts. Pathogen fitness in vaccinated hosts is typically reduced, and so increased pathogen transmissibility is selected for [11–17]. As increasing transmission often requires an increase in virulence [18–21], more virulent pathogens will be favoured. The evolution of higher virulence as a consequence of vaccination has been documented empirically in Marek’s disease [22–24], experimentally in rodent malaria [25, 26], and has been implicated as a contributing factor to the re-emergence of pertussis [27].

Although both the vaccine-induced evolution of escape [e.g., 28–32] and virulence [11–14, 32] are well-studied, these adaptations have typically been considered in isolation from one another. When they have been considered in combination [17], the focus was their long-term evolution and so the possibility of transient multilocus dynamics was ignored. Here we develop a model at the interface of epidemiology and population genetics to study both the short- and long-term multilocus adaptation of pathogens to vaccination. Our goal is to understand the role of (deterministic) selection and so we neglect the influence of stochasticity and chance mutations [e.g., 33–36]. We focus upon the protective effects of vaccination acting at different stages of the pathogen’s life-cycle, and show that these have important consequences for the sign of epistasis between virulence and vaccine-escape; this is key to understanding the evolutionary dynamics. We then examine the role of recombination in shaping the evolutionary outcome.

## Model

Consider a standard SIR model (Sup. Info. S1.1). Transmission of the pathogen occurs via mass-action with rate constant *β*, while pathogen infection causes virulence-related mortality at a per-capita rate *α*. A positive relationship between virulence and transmission, potentially mediated by within-host pathogen growth, has been observed for many infectious diseases [18, 19, 21, 37], including respiratory viruses affecting the lower respiratory tract [38]. To indicate the possibility of this relationship, we will write the rate constant *β* as *β*[*α*]. Infections are cleared at rate *γ*, and recovered hosts are fully protected from future infections through naturally-acquired immunity. Hosts enter the population at a fixed rate *λ* (e.g., through birth or immigration) and are removed due to natural causes (e.g., death or emigration) at a per-capita rate *d*.

A fraction *p* of the hosts entering the population are vaccinated. The vaccine has five protective effects that can alter distinct steps of the pathogens life-cycle (Fig. 1**a**):

1. it reduces the risk of infection by a factor 0 ≤ *ρ*_1_ ≤ 1;
2. it reduces within-host pathogen growth by a factor 0 ≤ *ρ*_2_ ≤ 1, affecting both virulence and transmission;
3. it reduces infectiousness (without affecting virulence) by a factor 0 ≤ *ρ*_3_ ≤ 1;
4. it reduces virulence (without affecting transmission) by a factor 0 ≤ *ρ*_4_ ≤ 1; and,
5. it reduces the duration of infection by increasing the recovery rate by a factor 0 ≤ *ρ*_5_.

**Figure 1:**
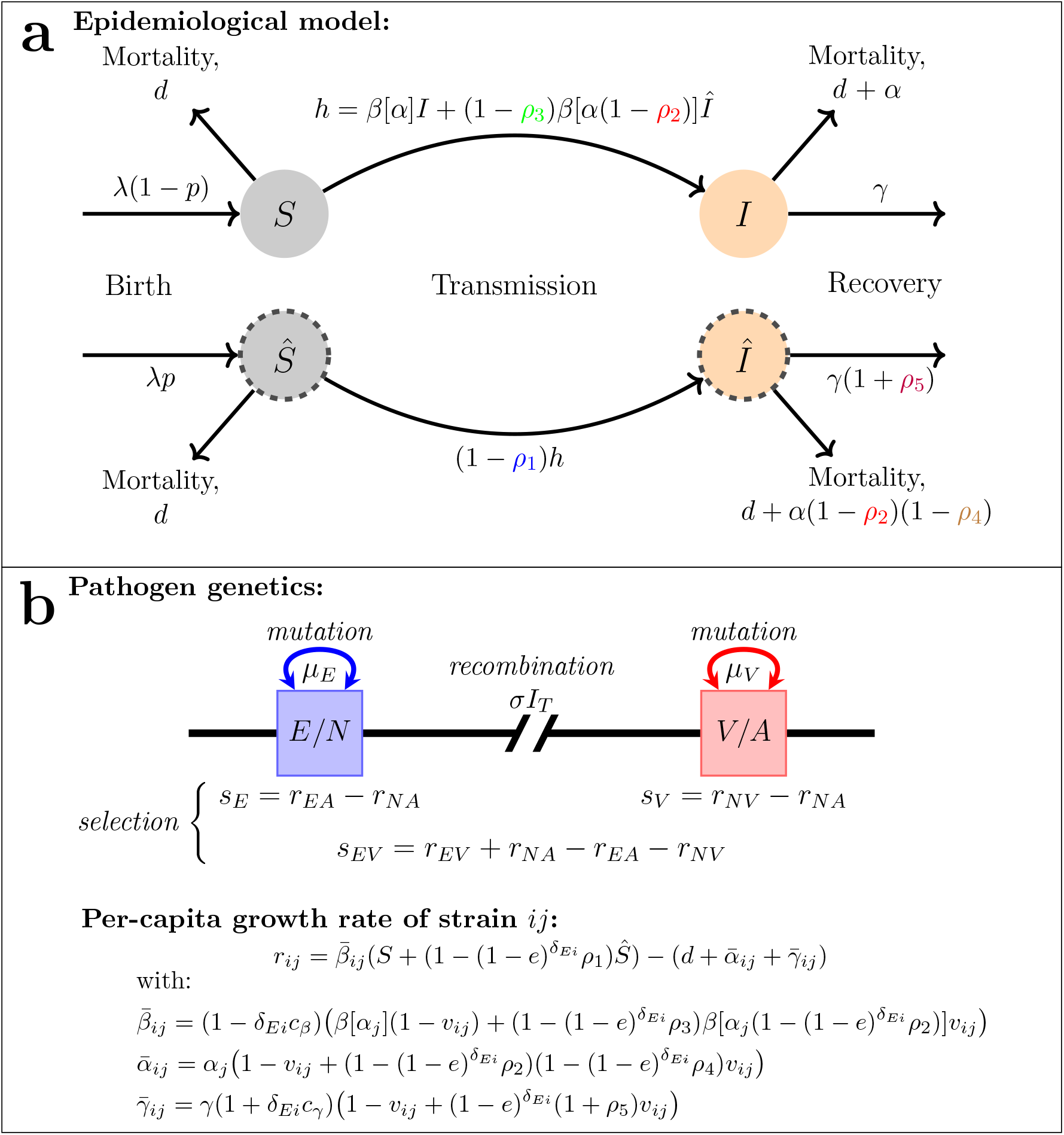
Schematic of the epidemiological model (panel a) and the evolutionary model (panel b). In panel **a**, we highlight how each of the possible vaccine protections affect the epidemiological dynamics. In particular, vaccines may: reduce the risk of infection (*ρ*_1_), reduce within-host pathogen growth (*ρ*_2_), reduce infectiousness (*ρ*_3_), reduce virulence (*ρ*_4_), and/or hasten infection clearance (*ρ*_5_). In panel **b**, we highlight the two-locus, diallelic evolutionary model. Each locus undergoes mutation (*μ*_*V*_, *μ*_*E*_), while recombination occurs between loci (*σI*_*T*_). Selection occurs through the additive selection coefficients (*s*_*E*_, *s*_*V*_), and epistasis (*s*_*EV*_); each of these is defined in terms of the per-capita growth rates of the different strains, *r*_*ij*_ [42, 50]. In turn, the per-capita growth rates, *r*_*ij*_, depend upon the epidemiological model (and so are not constant), and are the average growth rate of strain *ij* across the two selective environments, vaccinated and unvaccinated hosts. Note that *δ*_*Ei*_ is the Kronecker delta, and is equal to one if *E* = *i* and zero otherwise.

Vaccine protection is assumed to be lifelong. Let *S* and 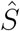 denote the density of unvaccinated and vaccinated susceptible hosts, and *I* and 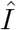 denote the density of unvaccinated and vaccinated infected hosts at time *t*.

The pathogen has two diallelic loci of interest (Fig. 1**b**). The first locus controls the pathogens ability to escape (allele *E*) or not (allele *N*) vaccine protection, by reducing each of the *ρ_i_* by a factor *e*; thus *e* = 0 corresponds to no vaccine-escape while *e* = 1 corresponds to complete escape. Carrying allele *E*, however, may come with fitness costs that reduce transmission by a factor 0 ≤ *c*_*β*_ ≤ 1, and/or increase rate of infection clearance by a factor *c*_*γ*_ ≥ 0. The second locus controls the pathogens virulence. Strains carrying allele *A* express virulence *α*_*A*_, whereas strains carrying allele *V* express virulence *α*_*V*_, and are more virulent and transmissible (i.e., *α*_*V*_ > *α*_*A*_ and *β*[*α*_*V*_] > *β*[*α*_*A*_]). There are thus four possible pathogen strains, *ij* ∈ {*NA, NV, EA, EV*}, and we will denote the per-capita growth of strain *ij* as *r*_*ij*_. Mutations occur at the escape and virulence locus at per-capita rates *μ*_*E*_ and *μ*_*V*_, respectively, while recombination (or genetic reassortment) between strains occurs at a per-capita rate *σI*_*T*_, where *I*_*T*_ is the total infection density and *σ* is a rate constant (Fig. 1**b**).

Let *f*_*ij*_ and *f*_*k*_ denote the frequency of strain *ij* and allele *k*, respectively, and *D* the linkage disequilibrium (LD) between alleles *E* and *V*, that is, *D* = *f*_*EV*_ − *f*_*E*_*f*_*V*_ [39]. Further, denote the additive selection coefficients for vaccine-escape and (increased) virulence as *s*_*E*_ ≡ *r*_*EA*_ − *r*_*NA*_ and *s*_*V*_ ≡ *r*_*NV*_ − *r*_*NA*_, respectively, and the epistasis in per-capita growth between alleles *E* and *V* as *s*_*EV*_ ≡ *r*_*EV*_ − *r*_*EA*_ + *r*_*NA*_ − *r*_*NV*_ (see Fig. 2). Then the evolutionary dynamics can be written

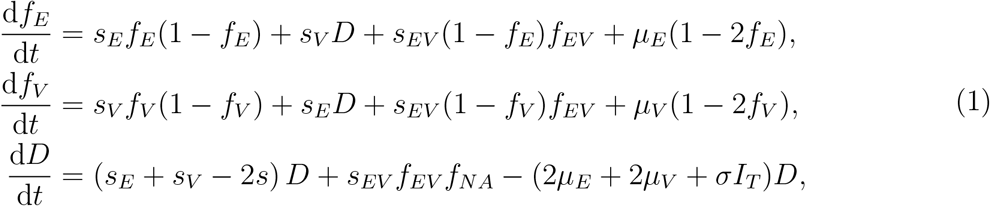

where *s* = *s*_*E*_*f*_*E*_ + *s*_*V*_ *f*_*V*_ + *s*_*EV*_ *f*_*EV*_. System (1) has the standard interpretation [e.g., 40–42]. The dynamics of the frequency of allele *E* (*mutatis mutandis* allele *V*) depend upon: (i) the action of direct selection, *s*_*E*_, proportional to the genetic variance, (ii) the influence of indirect selection, *s*_*V*_, mediated through LD, (iii) epistasis, *s*_*EV*_, and (iv) unbiased mutations. Thus the two key emergent quantities when considering the multilocus dynamics are the non-random assortment of alleles (LD; [39]) and any gene interactions (epistasis; [43]). Moreover, from (1), these two quantities are linked: LD is generated by epistasis (*s*_*EV*_ ≠ 0), which produces same sign LD [44, 45]. Once LD is present in the population, it can be amplified by directional selection, while it is removed by mutations and recombination [46, 47].

**Figure 2:**
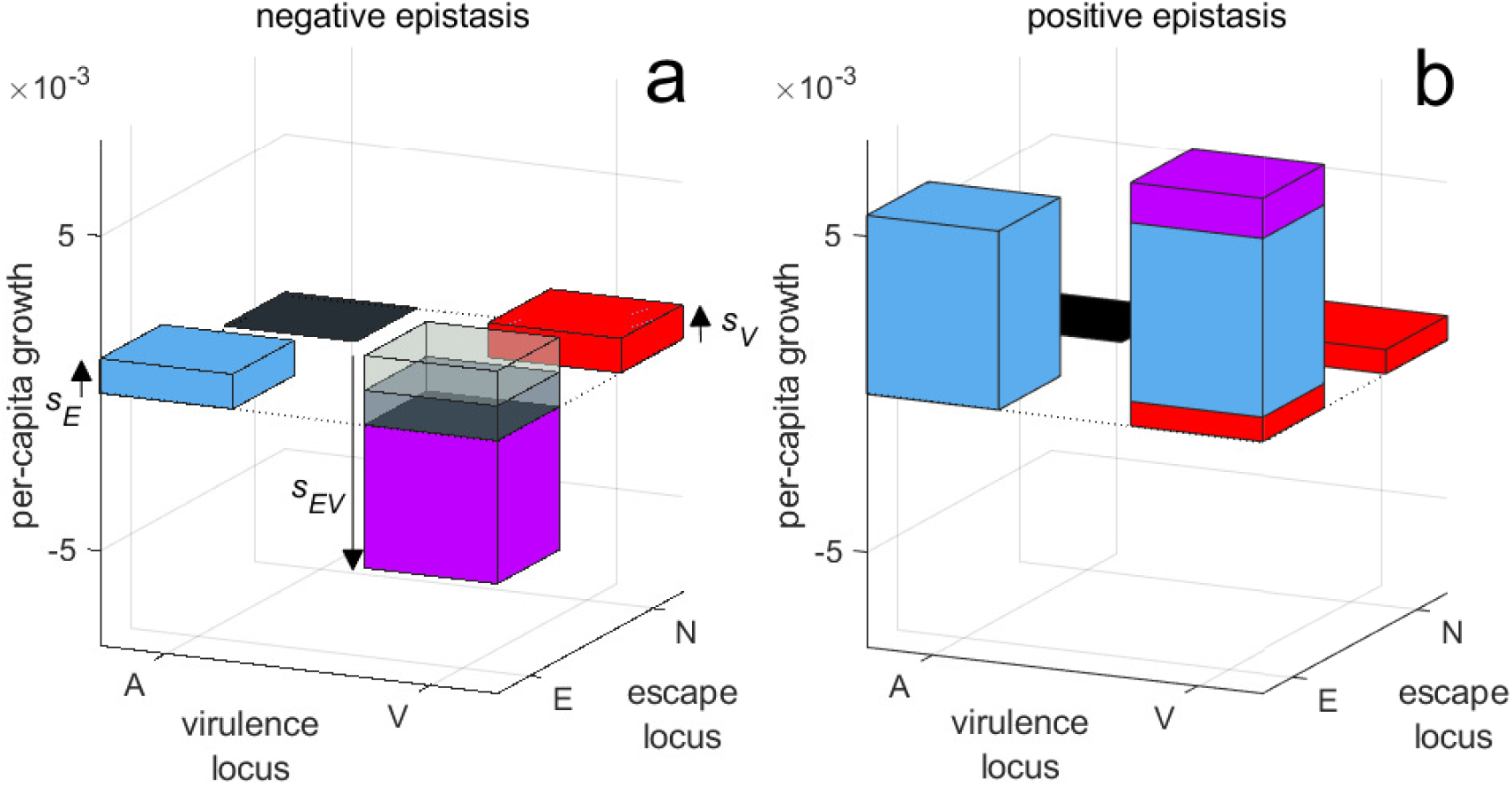
Fitness landscape at the strain *NA* equilibrium. Each bar represents the percapita growth rate of strain *ij* at the strain *NA* equilibrium (so *r*_*NA*_ = 0); the colours indicate the contribution of allele *E* (blue), allele *V* (red), and epistasis (magenta). Panel **a** shows an example of negative epistasis, which occurs when vaccines reduce virulence mortality (*ρ*_2_, *ρ*_4_), while panel **b** shows an example of positive epistasis, which occurs when vaccines reduce transmission (*ρ*_1_, *ρ*_3_) and increase clearance (*ρ*_5_). Parameter values used were *p* = 0.6, *e* = 1, 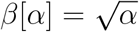, *λ* = *d* = 0.05, *γ* = 0.1; while for panel **a**, *ρ*_1_ = *ρ*_3_ = *ρ*_5_ = 0, *ρ*_2_ = 0.3, *ρ*_4_ = 0.01, *c*_*β*_ = 0, *c*_*γ*_ = 0.01, and for panel **b**, *ρ*_2_ = *ρ*_4_ = 0, *ρ*_1_ = 0.025, *ρ*_3_ = 0.02, *ρ*_5_ = 1, *c*_*β*_ = 0.03, *c*_*γ*_ = 0.45. Finally, *α*_*A*_ and *α*_*V*_ were chosen to be the optimal virulence on genetic background *N* in an entirely unvaccinated and vaccinated population, respectively (Sup. Info. S1.4).

In addition to the evolutionary dynamics described by (1), there is a system of differential equations describing the epidemiological dynamics. These equations can be found in Sup. Info. S1.2; here our focus is upon system (1). However, we note that because the selection coefficients and epistasis are defined in terms of the per-capita growth rates, *r*_*ij*_, which in turn depend upon the epidemiological dynamics, *s*_*E*_, *s*_*V*_ and *s*_*EV*_ can temporally vary in both magnitude and sign due to the epidemiological dynamics (see Fig. 1).

## Results

### Single locus evolution

First, consider evolution restricted to a single locus. This occurs if the mutation rate at one locus is much smaller than the other due to the absence of accessible mutations. For example, smallpox was unable to escape vaccine immunity, leading to its eradication [48, 49], while for other pathogens, evolutionary changes to virulence may not be possible due to various biological constraints.

If evolution is restricted to the escape locus (so *μ*_*V*_ = 0 and *f*_*V*_ = *D* = 0), system (1) reduces to

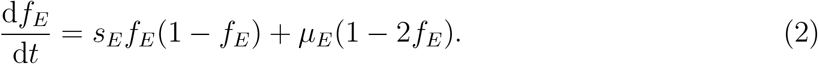

Thus the single locus dynamics are governed by the flux of mutation, *μ*_*E*_, and selection for the escape allele, *s*_*E*_. As mutation rate is expected to be small, the dynamics of *f*_*E*_ will be primarily shaped by selection: if *s*_*E*_ > 0, allele *E* is favoured. Hence from consideration of *s*_*E*_ (Fig. 1**b**), increasing the vaccine protection against infection (*ρ*_1_), growth (*ρ*_2_), transmission (*ρ*_3_) and clearance (*ρ*_5_), reduces strain *NA* fitness and so favours allele *E*. Likewise, decreasing the costs of escape increases transmissibility (*c*_*β*_) and duration of carriage (*c*_*γ*_) of strain *EA*, favouring allele *E* (Sup. Info. S1.3). Vaccine protection against virulence (*ρ*_4_) is an exception as it increases the duration of carriage without a concomitant reduction in transmission, and so increases strain *NA* fitness. An increasing degree of escape (increasing *e*) amplifies the protective effect(s) of vaccination against allele *N* relative to allele *E*, and so can either increasingly favour (*ρ*_1_, *ρ*_2_, *ρ*_3_, *ρ*_5_ > 0) or disfavour (*ρ*_4_ > 0) allele *E*. Generally speaking, if the costs of escape are not too high relative to the degree of escape, and vaccine protection and coverage are not too low, allele *E* is favoured over allele *N* on the genetic background *A* (*s*_*E*_ > 0).

If instead evolution is restricted to the virulence locus (so *μ*_*E*_ = 0 and *f*_*E*_ = *D* = 0), system (1) reduces to

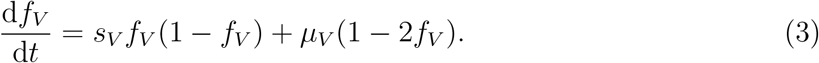

Thus the more virulent strain will be favoured by selection if *s*_*V*_ > 0. Selection for virulence on the genetic background *N* is a weighted average across two environments: unvaccinated and vaccinated hosts; the weights are the fraction of infections of a given strain that are found in either environment (Fig. 1**b**; Sup. Info. S1.4). However, depending upon how much allele *V* increases transmissibility relative to increasing virulence (as compared to allele *A*), on the genetic background *N* allele *V* can be unconditionally favoured even in an unvaccinated population, or unconditionally disfavoured even in an entirely vaccinated population. As vaccination plays a limited role in either case, suppose that allele *V* is neither unconditionally favoured nor disfavoured. Then if the vaccine affects virulence mortality (*ρ*_2_, *ρ*_4_) and/or clearance (*ρ*_5_), higher virulence and transmission will be selected for in vaccinated hosts to compensate for the reduced duration of infection [11–13, 15]. Whether higher virulence is favoured in the population as a whole, however, is determined by the balance between the two selective environments: increasing coverage (*p*) and decreasing protection from infection (*ρ*_1_) and transmission (*ρ*_3_), increases the importance of vaccinated hosts, favouring higher virulence [11–13, 15].

More broadly, the key competitive difference between alleles *E* and *N* is that infections carrying allele *N* are better at exploiting either unvaccinated hosts (strain *NA*) or vaccinated hosts (strain *NV*), but have reduced fitness in vaccinated hosts (strain *NA*) or unvaccinated hosts (strain *NV*), whereas infections carrying allele *E* pay a fixed cost to more equally exploit both types of hosts. Thus strain *Nj* is a *specialist* whose fitness depends upon the fraction of susceptible and infected hosts that are vaccinated, while in the extreme case where *e* = 1, strain *Ej* is a *generalist* whose fitness depends upon the availability of hosts.

### Epistasis and the production of linkage disequilibrium

Next, suppose that segregation is occurring at both loci, that is, both adaptations are mutationally accessible (*μ*_*E*_ > 0 and *μ*_*V*_ > 0). The key emergent quantity when considering multilocus dynamics is the non-random assortment of alleles (LD; [39]). From (1), LD is produced by epistasis, *s*_*EV*_; moreover, epistasis generates same sign LD [44, 45]. Epistasis occurs whenever the fitness of an allele depends upon its genetic background; in our model, this happens in two ways. The first is that vaccination creates two selective environments, causing allele *V* to be disproportionately affected by vaccination as compared to allele *A*, particularly on the genetic background *N*. The logic is straightforward: if the vaccine reduces virulence (*ρ*_2_, *ρ*_4_) or increases clearance (*ρ*_5_), higher virulence will disproportionately reduce the expected duration of infection in unvaccinated hosts relative to vaccinated hosts. Consequently, a greater proportion of strain *iV* infections will be found in vaccinated hosts than strain *iA* infections (Sup. Info. S1.2), while since escape reduces the effects of vaccination, this difference will typically be magnified on the genetic background *N* relative to *E*. The second way epistasis is generated is through any non-additive interactions between parameters associated with the virulence locus (*β*, *α*), and those associated with the escape locus (vaccine protections, costs of escape; [42]). Because *α*_*V*_ > *α*_*A*_ and *β*[*α*_*V*_] > *β*[*α*_*A*_], any such interactions will disproportionately affect infections carrying allele *V* over *A* for a given genetic background (*E* or *N*).

Next, what is the sign of the epistatic contribution of the different types of vaccine protection? Because escape reduces the impact of vaccination, manipulating the vaccine protections will typically disproportionately affect the difference in per-capita growth rate (fitness) between strains *NA* and *NV* as compared to strains *EA* and *EV*. Since epistasis is defined as *s*_*EV*_ ≡ *r*_*EV*_ − *r*_*EA*_ + *r*_*NA*_ − *r*_*NV*_ [42, 50], and as the difference *r*_*NA*_ − *r*_*NV*_ will be larger in magnitude than the difference *r*_*EA*_ − *r*_*EV*_, to predict the sign of the epistatic contribution of a given vaccine protection, assuming escape is not to ineffective (*e* not too small; see Sup. Info. S1.5), it suffices to consider whether interactions between vaccine protections and virulence will disproportionately increase or decrease the fitness of strain *NA* relative to strain *NV*. If strain *NA* experiences an increase (resp. decrease) in fitness relative to strain *NV*, then positive (resp. negative) epistasis will be produced (Sup. Info. S1.5). With this in mind, the predictions are as follows:

1. **Vaccines that reduce transmission** (*ρ*_1_, *ρ*_2_, *ρ*_3_) **produce positive epistasis**. Since allele *V* is more transmissible than allele *A*, and as a larger fraction of *NV*-infections are in vaccinated hosts than *NA*-infections, on the genetic background *N* allele *V* will experience a disproportionately higher reduction in transmission due to vaccination than allele *A*. Thus strain *NV* is at a disadvantage relative to strain *NA*, and so vaccines reducing transmission produce positive epistasis.
2. **Vaccines that reduce virulence mortality** (*ρ*_2_, *ρ*_4_) **produce negative epistasis**. Ignoring any concomitant effects to transmission, reducing virulence mortality increases pathogen fitness. Since allele *V* corresponds to more virulent infections, and as a larger fraction of *NV*-infections are in vaccinated hosts as compared to *NA*-infections, allele *V* will disproportionately benefit from any reductions in virulence. Thus strain *NV* will be at an advantage relative to strain *NA*, and so vaccines reducing virulence mortality produce negative epistasis.
3. **Vaccines that increase clearance** (*ρ*_5_) **produce positive epistasis**. Because a larger fraction of *NV*-infections are in vaccinated hosts than *NA*-infections, on the genetic background *N* allele *V* will be disproportionately negatively affected by increased clearance due to vaccination. Consequently, strain *NV* will be at a disadvantage relative to strain *NA* and so vaccines increasing clearance produce positive epistasis.

Note that vaccines which block growth (*ρ*_2_) have two opposing contributions: by reducing transmission, *ρ*_2_ produces positive epistasis, and by reducing virulence mortality, *ρ*_2_ produces negative epistasis. Thus depending upon the epidemiology, as well as the difference between *α*_*A*_ and *α*_*V*_ and/or *β*[*α*_*A*_] and *β*[*α*_*V*_], the epistatic contribution of *ρ*_2_ can be positive or negative. More generally, the relative importance of the different factors contributing to epistasis can temporally vary; for example, if susceptible hosts are abundant, the contribution of transmission to epistasis is likely to be more important (Sup. Info. S1.5).

### Negative epistasis and evolutionary bistability

What are the dynamical implications of the sign of epistasis? First, suppose the vaccine protections produce negative epistasis (i.e., vaccines reduce virulence mortality, *ρ*_2_, *ρ*_4_; Fig. 2**a**), and there is directional selection favouring alleles *E* and *V* (*s*_*E*_ > 0, *s*_*V*_ > 0) at the strain *NA* endemic equilibrium. In this circumstance, strains *EA* and *NV* will initially increase, at a rate dictated by the strength of selection for allele *E* and *V*, respectively. As negative epistasis disfavours strain *EV*, the increase in strains *EA* and *NV* will cause an increase in negative LD until alleles *E* and *V* are sufficiently abundant such that *f*_*E*_ + *f*_*V*_ ≈ 1. At this point, and assuming mutations and recombination are sufficiently infrequent, system (1) reduces to

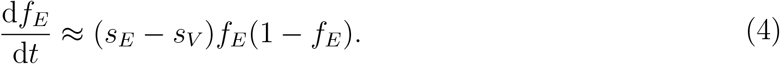

From (4), epistasis no longer directly impacts the dynamics of allele frequencies. Instead whichever of alleles *E* or *V* is more strongly selected for (larger selection coefficient) will increase in frequency at the expense of the other; that is, strains *EA* and *NV* are in direct competition with each other. This competition will cause the population to evolve along the curve in (*f*_*E*_, *f*_*V*_, *D*)-space such that *f*_*V*_ = 1 − *f*_*E*_ and *D* = − *f*_*E*_(1 − *f*_*E*_) (denoted Γ(*f*_*E*_) in Fig. 3) according to equation (4), until either strain *EA* or strain *NV* outcompetes the other (Fig. 3).

**Figure 3:**
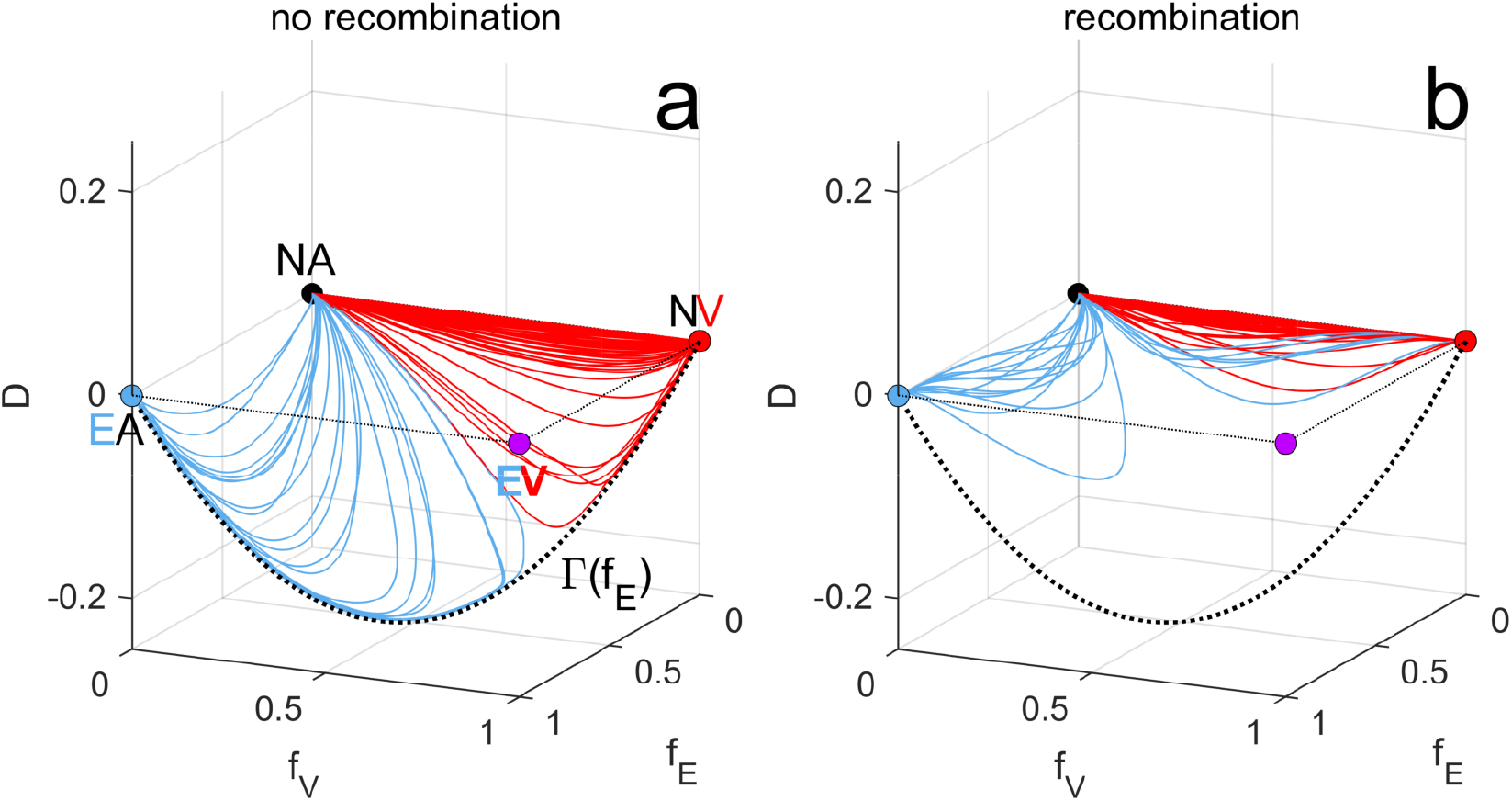
Negative epistasis and recombination can lead to evolutionary bistability. If both vaccine-escape and virulence are advantageous (*s*_*E*_ > 0 and *s*_*V*_ > 0), but epistasis is negative (*s*_*EV*_ < 0), alleles *E* and *V* are in competition. Starting from the strain *NA* equilibrium (solid black circle), strains *NV* and *EA* will increase in frequency, producing negative LD until the population is in the vicinity of the curve, Γ(*f*_*E*_) (dashed black curve; see Sup. Info. S1.6). Along Γ(*f*_*E*_), strains *EA* and *NV* are in direct competition, and so whichever allele (*E* or *V*) has the larger selection coefficient will go to fixation (panel **a**, *σ* = 0); blue curves correspond to fixation of strain *EA*, red curves to fixation of strain *NV*. Recombination can create an evolutionary bistability such that ‘faster’ growing strains tend to reach fixation, even if they are competitively inferior (panel **b**, *σ* = 0.05); here the colours indicate which strain would fix in the absence of recombination. Each simulation starts with a monomorphic pathogen population (only the *NA* genotype is present initially) at it’s endemic equilibrium following vaccination; mutation introduces genetic variation and allows pathogen adaptation to vaccination. Panels **a**,**b** show 100 simulations using the parameter set: *p* = 0.6, *e* = 1, 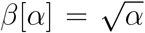, *λ* = *d* = 0.05, *γ* = 0.1, *μ*_*E*_ = 10^−4^, *μ*_*V*_ = 10^−4^, *ρ*_1_ = *ρ*_3_ = *ρ*_5_ = 0, and with *ρ*_2_, *ρ*_4_, *c*_*β*_, *c*_*γ*_ chosen uniformly at random on the intervals [0, 0.95], [0, 0.95], [0, 0.05], [0, 0.25], respectively, subject to the constraints that at the strain *NA* equilibrium, *s*_*E*_ > 0, *s*_*V*_ > 0, and *α*_*V*_/*α*_*A*_ > 1.25. *α*_*A*_ and *α*_*V*_ were chosen to be the optimal virulence in an entirely unvaccinated and vaccinated population, respectively (Sup. Info. S1.4).

Recombination affects these dynamics: by removing LD from the population, recombination prevents LD from increasing in magnitude [46, 47]. Consequently, the combination of strain competition and recombination mean that it no longer need be true that the ‘fitter’ strain ultimately reaches fixation if doing so requires the population to first pass through an intermediate state with a large amount of LD. As such, recombination can create a bistability between the *NV* - and *EA*-strain equilibria (Fig. 3**b**); in fact, in the limiting case of quasi-linkage equilibrium [51–53], recombination greatly increases the likelihood that the *NV* - and *EA*-equilibria are simultaneously locally stable attractors (Sup. Info. S1.8). The evolutionary implications of this bistability is that historical contingency, or being ‘first’, is more important than being ‘fitter’. Specifically, if one allele (either *E* or *V*) is more mutationally accessible, or is associated with more rapid growth from the strain *NA* equilibrium (i.e., has a larger selection coefficient), even if this allele is competitively inferior to the other in the long-term (in the absence of recombination), it is likely to reach fixation and exclude the other (Fig. 3**a**,**b**).

### Positive epistasis and the evolution of virulence

Next, suppose epistasis is positive (i.e., vaccines reduce infection/transmission, *ρ*_1_, *ρ*_3_, and/or increase clearance, *ρ*_5_; Fig. 2**b**), and *s*_*E*_ > 0 and *s*_*V*_ > 0 at the strain *NA* equilibrium. When does strain *EV* (and so positive LD) transiently increase in the population? If the selection coefficients are of comparable magnitude, *s*_*E*_ ≈ *s*_*V*_, alleles *E* and *V* increase in the population at a similar speed, and so positive epistasis acts to transiently favour strain *EV* (and positive LD; Fig. 4**a**). On the other hand, if one selection coefficient is much larger than the other (say *s*_*E*_ ≫ *s*_*V*_), then allele *E* will rapidly increase, favouring whichever virulence allele it is initially associated with (i.e., the virulence allele hitch-hikes [54]). Consequently, if mutations at each locus are independent, strain *EA* will transiently dominate as it is more mutationally accessible from strain *NA* (Fig. 4**a**). If instead double mutations (e.g., *NA* mutates to *EV*) occur with comparable frequency to single mutations, or strains *EA* and *EV* are initially equally abundant (e.g., due to chance fluctuations), strain *EV* (and positive LD) will transiently dominate, as it benefits from both the positive epistasis and selection on allele *V*. One key determinant of the relative magnitudes of *s*_*E*_ and *s*_*V*_ will be the costs (*c*_*β*_, *c*_*γ*_), and degree (*e*), of vaccine escape. If costs are low and vaccine escape is easily accessible, typically *s*_*E*_ ≫ *s*_*V*_, whereas costly and/or limited escape can lead to *s*_*V*_ ≫ *s*_*E*_.

**Figure 4:**
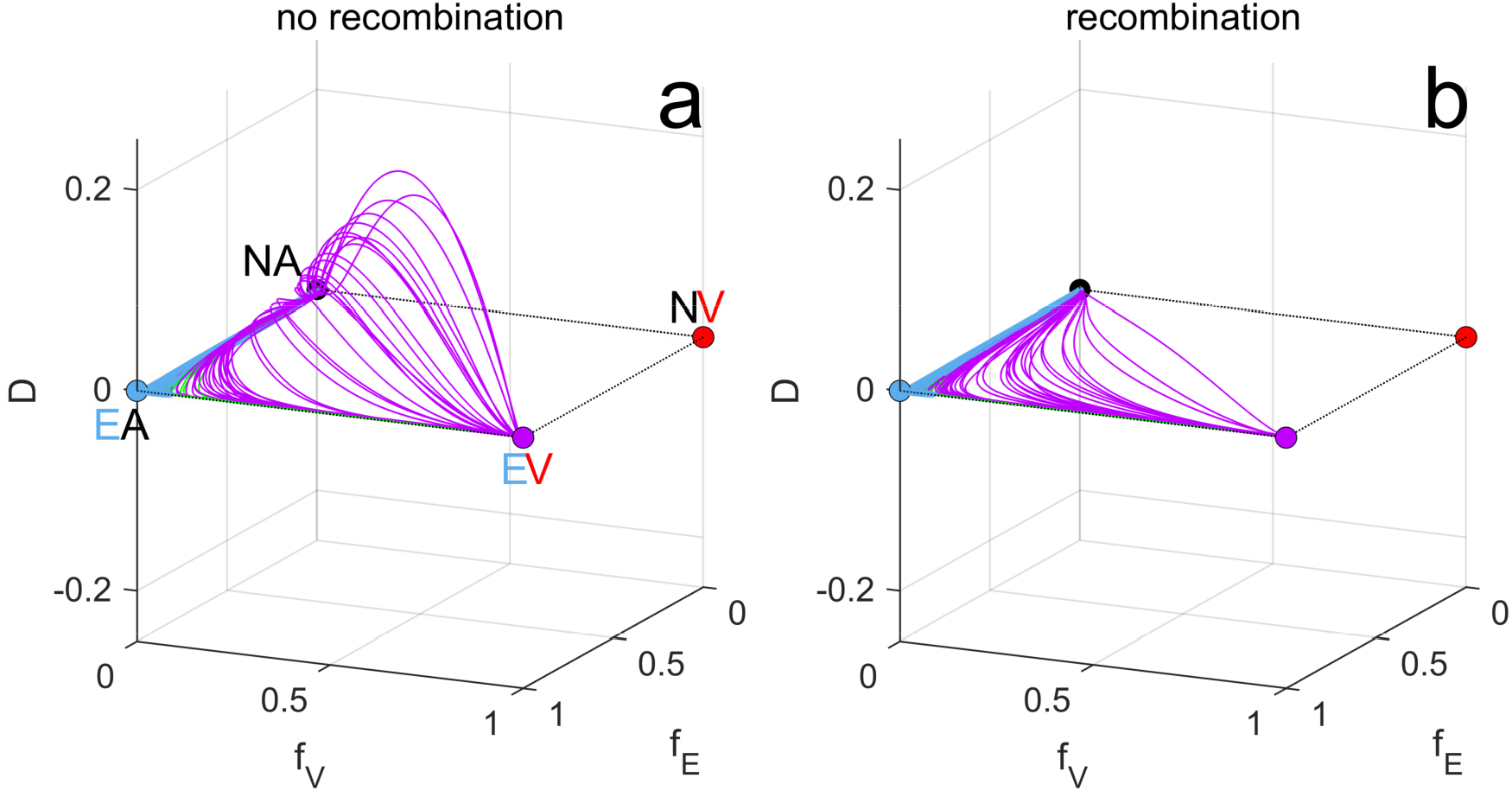
Positive epistasis and the evolution of virulence. Positive epistasis favours strains that are both more virulent and evasive in the short− and long-term leading to the build up of LD (panel **a**; *σ* = 0). Frequent recombination breaks up LD (panel **b**; *σ* = 0.05), favouring the sequential fixation of traits; in this case, allele *E* reaches quasi-fixation first followed by allele *V*. Colours indicate which strain fixes: blue is strain *EA*, magenta is strain *EV*. Each simulation starts with a monomorphic pathogen population (only the *NA* geno-type is present initially; solid black circle) at it’s endemic equilibrium following vaccination; mutation introduces genetic variation and allows pathogen adaptation to vaccination. Panels **a**,**b** show 100 simulations using the parameter set: *p* = 0.6, *e* = 1, 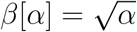, *λ* = *d* = 0.05, *γ* = 0.1, *μ*_*E*_ = 10^−4^, *μ*_*V*_ = 10^−4^, *ρ*_2_ = *ρ*_4_ = *c*_*β*_ = 0, and with *ρ*_1_, *ρ*_3_, *ρ*_5_, *c*_*γ*_ chosen uniformly at random on the intervals [0, 0.9], [0, 0.9], [0, 10], [0, 5], respectively, subject to the constraints that at the strain *NA* equilibrium, *s*_*E*_ > 0, *s*_*V*_ > 0, and *α*_*V*_/*α*_*A*_ > 1.25. *α*_*A*_ and *α*_*V*_ were chosen to be the optimal virulence in an entirely unvaccinated and vaccinated population, respectively (Sup. Info. S1.4).

In the long-term, one (or both) of the alleles will approach quasi-fixation and LD will vanish. Suppose allele *E* reaches quasi-fixation; then the dynamics of (1) reduce to

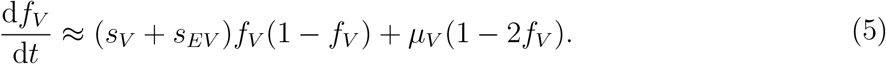

Here *s*_*V*_ + *s*_*EV*_ = *r*_*EV*_ − *r*_*EA*_ is the selection coefficient for allele *V* on the genetic background *E*; thus whether allele *A* or *V* is ultimately favoured depends upon its sign. For example, suppose escape is near complete (*e* ≈ 1) and costs of escape to transmission are negligible (*c*_*β*_ 0). Then the sign of *s*_*V*_ + *s*_*EV*_ is determined by the available hosts: if hosts are abundant, more transmissible strains are favoured [19, 55–57]. One factor affecting the available hosts is the costs of escape to duration of carriage (*c*_*γ*_). Increasing *c*_*γ*_ reduces total infections, thereby increasing hosts and favouring allele *V* (Sup. Info. S1.7).

Recombination can affect these dynamics. By breaking up LD, recombination can prevent the transient selection for the *EV* strain. If instead strain *EV* is favoured both transiently and in the long-term, frequent recombination will prevent positive LD from building up in the population. Therefore evolution occurs sequentially such that either allele *E* or *V* will fix first (depending upon which allele is more strongly selected for), before evolution proceeds at the other loci (Fig. 4**b**).

## Discussion

Unlike antimicrobials, vaccines typically provide evolutionarily robust protection, and have had some remarkable successes [1–3]. However, although less common than antimicrobial resistance, pathogen evolution in response to vaccination does occur [1, 3, 24, 58–60]. Two primary adaptive routes have been identified along which pathogens adapt to vaccination [24]: by evading the vaccine-induced immune response, or by adjusting life history traits, specifically virulence and transmission. Although each of these adaptations have been well-studied in isolation [e.g., 11–14, 28, 30, 31], their simultaneous evolution is poorly understood. Here we considered the joint evolution of vaccine-escape and virulence, with a focus upon the role played by the different possible protections conferred by vaccination. We considered three classes of vaccine protections: (i) those that reduce pathogen transmission either by blocking infection (sterilizing immunity) or reducing infectiousness [61, 62]; (ii) those that increase clearance rate (e.g., by inducing a T-cell response [63]); and (iii) those that reduce virulence (e.g., by blocking growth or removing virulence factors [64]). Vaccines reducing infectivity, transmission and/or increasing clearance generate positive epistasis, and so strains that carry both vaccine-escape and virulence mutations can be favoured transiently and in the long-term. In contrast, vaccines reducing virulence mortality generate negative epistasis, favouring one adaptative route or the other, but not both.

From a public health standpoint, the least desirable situation is the evolution of strains that escape vaccine immunity and are more virulent; this occurs when epistasis is positive. In this case, the dynamics are very sensitive to the initial conditions, and significant positive LD (strain *EV* over-representation) can build-up transiently (Sup. Info. Fig. S3). This suggests stochasticity and chance are far more likely to play important evolutionary roles if epistasis is positive. By breaking up LD, recombination can prevent the transient evolution of more virulent, vaccine-escape strains, while if such strains are favoured in the long-term, recombination leads to the sequential fixation of mutations. Often, vaccine-escape mutations will reach quasi-fixation first, followed by the evolution of higher virulence. In this circumstance, if vaccines can be ‘tweaked’ to restore coverage once their efficacy begins to wane due to vaccine-escape (e.g., as for seasonal influenza), this will force the pathogen to continually evolve vaccine-escape, rather than becoming more virulent. This would suggest that proactive updates to vaccines may be advisable; although this may be impractical with conventional technology (e.g., updating/producing seasonal influenza vaccines takes months [65]), recent technological advances allow vaccines to be rapidly updated and manufactured (e.g., mRNA vaccines [66]).

The competition between vaccine-escape and virulence (negative epistasis) may have implications for ‘universal’ vaccines [8–10, 67]. Universal vaccines target more conserved viral components than ‘conventional’ vaccines, and so are expected to provide broader cross-protection against a range of pathogen subtypes, thereby preventing and/or slowing antigenic escape [8, 9]. However, the evolutionary pressures imposed by universal vaccines are not fully understood [10]. To date, models have primarily focused upon the potential of universal vaccines to reduce antigenic escape [8, 9], and have neglected other potential adaptive responses (e.g., virulence evolution). Reducing the evolutionary likelihood of vaccine-escape (by reducing *μ*_*E*_) could induce collateral evolutionary damage by tilting the competitive balance in favour of virulence (assuming *μ*_*V*_ is also not small). Importantly, the consequences of virulence evolution will depend upon the mechanism of protection offered by universal vaccines. For example, in influenza there are two broad categories of universal vaccine candidates: those that target haemagglutinin proteins [68, 69] and those that induce a broadly protective T-cell immune response [8, 10, 68]. Vaccines in the former category are expected to block infection [68], while vaccines in the latter are not, and instead are likely to reduce growth, disease severity, and duration of infection [8, 10, 70]. Although both categories of vaccines should reduce the likelihood of escape [8, 9], our analysis suggests that by only weakly blocking infection and by decreasing disease severity, vaccines inducing a broadly protective T-cell immune response will also favour selection for increased virulence. More broadly, understanding the evolutionary response to universal vaccines should not focus solely upon antigenic drift.

The sequencing of SARS-CoV-2 is unveiling the complex evolutionary dynamics taking place at multiple sites in the virus genome. Current attempts to understand the dynamics of the multitude of SARS-CoV-2 *variants* is overwhelming. Yet, different variants often share the same mutations affecting, for example, transmission (e.g., D614G [71, 72]) or vaccine-escape (e.g., E484K [73, 74]). Hence, studying the dynamics of the mutations, rather than the variants, can provide greater insight. For example, as recombination is common in coronaviruses [75], including SARS-CoV-2 [e.g., 76–78], one pressing question is whether it will produce a variant that is both highly transmissible/virulent and exhibits vaccine-escape [76, 79]. Although this is a worrisome possibility, our *mutation*-centered analysis emphasizes that such a variant need not be ‘fitter’: depending upon the vaccine protections, these two adaptations (mutations) may produce negative epistasis and so be disfavoured in combination. Indeed, the SARS-CoV-2 vaccines provide multifaceted protection, reducing infection [80, 81], infectiousness [61, 62], virulence [81] and infection duration [82]. The relative strengths of these protections will determine the sign, and magnitude, of epistasis, and it is epistasis that determines the evolutionary fate of recombinant variants, not the fitness of the ‘parental’ variants. Furthermore, in addition to combining different mutations, recombination also acts to break-up LD between mutations. As the rate of recombination depends upon the likelihood of co-infection (and so the density of infections), non-pharmaceutical interventions (NPIs) such as social-distancing, travel-restrictions, or mask-wearing, will reduce its rate. Previous work suggests that the usage of NPIs in response to the SARS-CoV-2 pandemic may affect the epidemiological dynamics of seasonal influenza and respiratory syncytial virus [83]; our work indicates that limiting recombination may have other evolutionary consequences, beyond restricting the generation of novel variants.

More generally, the framework we have developed here could equally be applied to other situations involving multiple interacting loci, such as other life history trade-offs (e.g., virulence-clearance [11, 19, 84] or clearance-transmission [85]), distinct loci controlling different aspects of immune-escape (e.g., antibody vs T-cell evasion), or compensatory mutations. Multilocus models constitute a particularly relevant framework to account for the joint evolution of multiple mutations and to provide an complementary perspective to the variant-centered view that is currently being used to understand the ongoing pandemic of SARS-CoV-2.

## S1 Supplementary Information

### S1.1 Model

Consider a standard SIR model. Transmission of the pathogen occurs via mass-action with rate constant *β*, while infection causes virulence-related mortality at a per-capita rate *α*. We allow for the possibility of a positive relationship between virulence and transmissibility, mediated through within-host replication rate [20, 21], and we indicate the potential functional dependence of transmission upon virulence by writing *β* as *β*[*α*]. Infections are naturally cleared at rate *γ*; following clearance, hosts have long-lasting naturally-acquired immunity. Hosts enter the population at rate *λ* (e.g., birth/immigration), and are removed from the population at a per-capita rate *d* (e.g., death/emigration). A fraction *p* of the hosts entering the population are vaccinated. Vaccination has five possible protective effects:

1. It reduces the probability of infection by a factor 0 ≤ *ρ*_1_ ≤ 1.
2. It reduces the growth of the pathogen within the host, and so reduces virulence by a factor 0 *ρ*_2_ 1. Since transmission is a function of virulence, this will also reduce transmission.
3. It reduces transmissibility (without affecting virulence) by a factor 0 ≤ *ρ*_3_ ≤ 1.
4. It reduces pathogen virulence (without affecting transmission) by a factor 0 ≤ *ρ*_4_ ≤ 1.
5. It hastens the clearance of the pathogen by a factor 0 ≤ *ρ*_5_. Vaccine protection is life-long.

The pathogen has two diallelic loci of interest. The first locus controls vaccine escape: infections carrying the escape allele, *E*, can evade vaccine protection, while infections with the non-escape allele, *N*, cannot. Specifically, if an infection carries the escape allele, then each of the protective effects of vaccination are reduced by a factor *e*; thus if *e* = 0, there is no evasion, whereas if *e* = 1, escape is complete. Carriage of the escape allele comes at the cost of reduced transmissibility by a factor 0 ≤ *c*_*β*_ ≤ 1, that is, (1 − *c*_*β*_)*β*[*α*], and/or an increased rate of clearance by a factor *c_γ_* ≥ 0, that is, *γ*(1 + *c_γ_*). The second locus controls the virulence of an infection, and we suppose there are two virulence alleles, *A* and *V*, corresponding to virulence *α_A_* and *α_V_*, respectively. On the same genetic background, infections carrying allele *A* are of lower virulence and are less transmissible than infections carrying allele *V*, that is, *α_V_ > α_A_* and *β*[*α_V_*] > *β*[*α_A_*]. In general, we will focus upon situations in which, on the genetic background *N*, allele *A* is ‘fitter’ in unvaccinated hosts, while allele *V* is ‘fitter’ in vaccinated hosts; we will discuss this in more detail later (Sup. Info. S1.4).

There are thus four pathogen strains, *ij* ∈ {*NA, NV, EA, EV*}, and we will let *I*_*ij*_ and 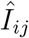 denote the density of *ij*-infections in unvaccinated and vaccinated hosts, respectively, at time *t*. Mutations occur at the escape and virulence loci at per-capita rates *μ_E_* and *μ*_*V*_, respectively. Recombination (or genetic reassortment) between the two loci may also occur. Specifically, if *σ* is a rate constant controlling the incidence of recombination, then recombination between strains *ij* and *kl* occurs at rate 2*σI*_*ij*_ *I*_*kl*_. Given recombination has occurred, either strain is equally likely to be the ‘recipient’ or ‘donor’ of the allele at a loci chosen with equal probability. Thus the change in density of *ij* infections in unvaccinated and vaccinated hosts, respectively, due to recombination is

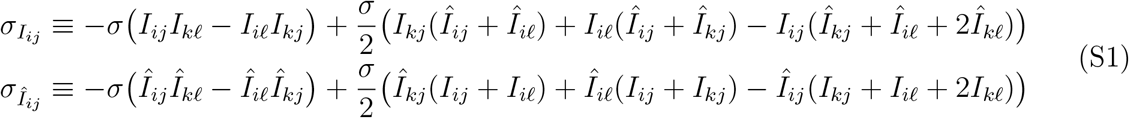

where *i* ≠ *k* and *j* ≠ *l*.

Let *S* and 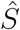 denote the density of uninfected unvaccinated and vaccinated hosts, respectively, and let *h*_*ij*_ denote the force of infection of strain *ij*, that is,

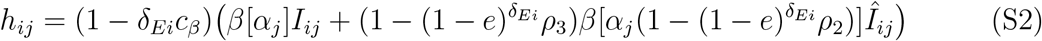

where *δ*_*Ei*_ is the Kronecker delta, equal to one if *E* = *i* and zero otherwise. Then the dynamics of the different strains in vaccinated and unvaccinated hosts are

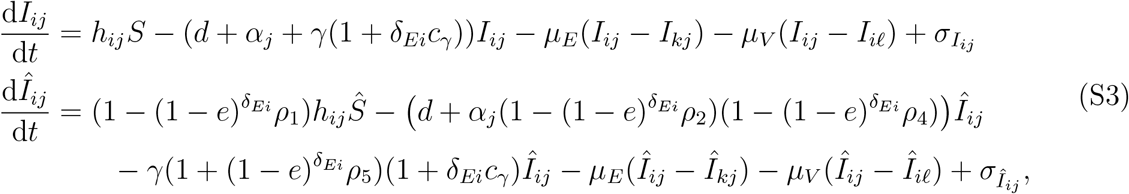

for *i* ≠ *k* and *j* ≠ *l*. Likewise, the dynamics of *S* and 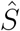 are

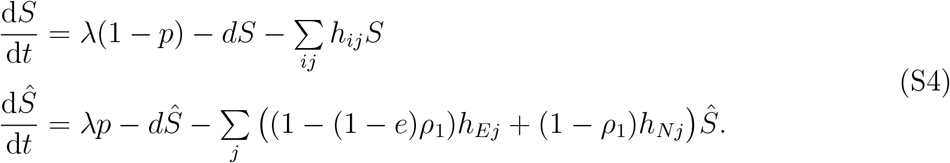

From the preceding assumptions, if we let 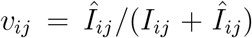 denote the fraction of *ij*-infections found in vaccinated hosts, the per-capita growth rate of *ij*-infections (so ignoring mutation and recombination), denoted *r*_*ij*_, is equal to

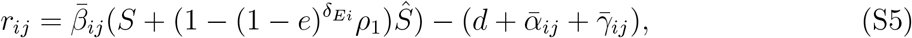

where

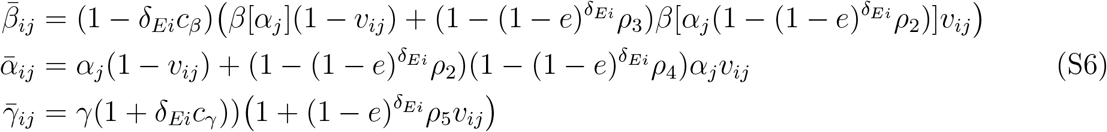

denote the average transmission, virulence, and clearance, respectively, of strain *ij*. The formulation of the *r*_*ij*_, specifically the 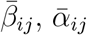, and 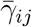 terms, emphasizes that the per-capita growth rate of *ij*-infections is a weighted average across selective environments (vaccinated and unvaccinated hosts), where the weights are the fraction of *ij*-infections in unvaccinated and vaccinated hosts, 1 − *v*_*ij*_ and *v*_*ij*_, respectively. If vaccine escape is complete (i.e., *e* = 1), then the per-capita growth rate of strain *Ej* will be unaffected by the distribution of *Ej*-infections across vaccinated and unvaccinated hosts.

### S1.2 Linkage disequilibrium formulation

We are interested in converting systems (S3) and (S4) into a form that separates the epidemiological. and evolutionary process. To do so let 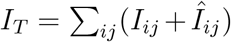 denote the total density of infections in the population, and let 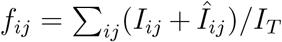 denote the frequency of strain *ij*-infections; it follows that *f*_*E*_ = *f*_*EA*_ + *f*_*EV*_ and *f*_*V*_ = *f*_*EV*_ + *f*_*NV*_ are the frequency of alleles *E* and *V*, respectively. Moreover, define the linkage disequilibrium (LD), or non-random association of alleles *E* and *V* [39], as

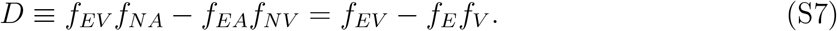

The additive selection coefficients for alleles *E* and *V* are defined as *s*_*E*_ ≡ *r*_*EA*_ − *r*_*NA*_ and *s*_*V*_ ≡ *r*_*NV*_ − *r*_*NA*_, respectively, while the epistasis in per-capita growth for alleles *E* and *V*, is defined as *s*_*EV*_ ≡ *r*_*EV*_ − *r*_*EA*_ + *r*_*NA*_ − *r*_*NV*_ (Fig. 2). Using this notation, the evolutionary variables are (*f*_*E*_, *f*_*V*_, *D*), and the evolutionary dynamics are described by

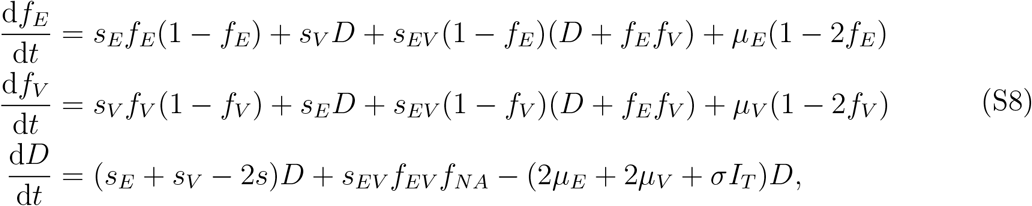

where *s* = *s*_*E*_*f*_*E*_ + *s*_*V*_ *f*_*V*_ + *s*_*EV*_ *f*_*EV*_.

In addition to the evolutionary dynamics we also have the set of epidemiological variables, (*S*, 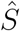, *I*_*T*_, *v*_*EA*_, *v*_*EV*_, *v*_*NA*_, *v*_*NV*_). The dynamical equations for susceptible hosts, *S* and 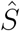 are provided in (S4). The other epidemiological variables describe the total density of infections, *I*_*T*_, and the fraction of *ij*-infections in vaccinated hosts, *v*_*ij*_. The equations describing these dynamics are

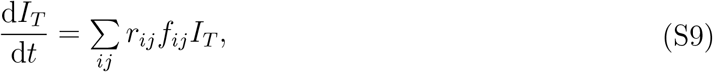

and

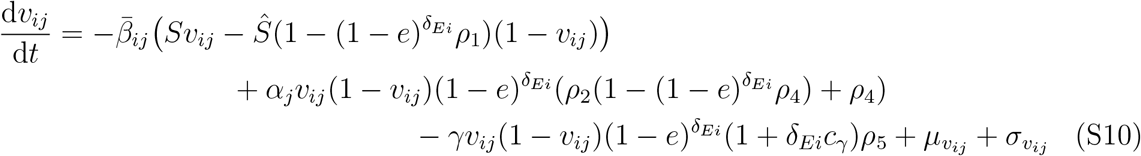

where

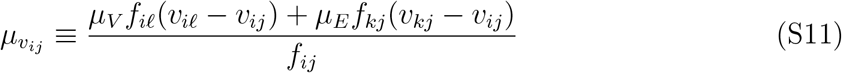

and

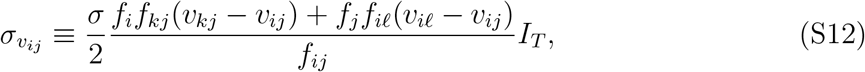

are the change in *v*_*ij*_ infections due to mutation and recombination, respectively.

#### S1.2.1 Relationships between the *v*_*ij*_(*t*)

In general, following any brief transient dynamics which may be under the influence of the initial conditions, we expect that *v*_*NV*_ ≥ (*t*) *v*_*NA*_(*t*). The logic is that strain *NA* is fitter in unvaccinated hosts, while strain *NV* is fitter in vaccinated hosts (see Sup. Info. S1.4). Numerical results suggest the relation *v*_*NV*_ (*t*) ≥ *v*_*NA*_(*t*) holds (Fig. S1); but to provide a rough mathematical argument for why, observe that *Nj* infections in unvaccinated hosts are produced at rate *h*_*Nj*_*S*, and on average will last (*d* + *α*_*j*_ + *γ*)^−1^ time units. Similarly, *Nj* infections in vaccinated hosts are produced at rate 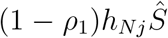, and on average will last (*d* + *α*_*j*_(1 − *ρ*_2_)(1 − *ρ*_4_) + *γ*(1 + *ρ*_5_))^−1^ time units. Thus we may be motivated to approximate *v*_*Nj*_(*t*) as 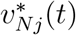, where

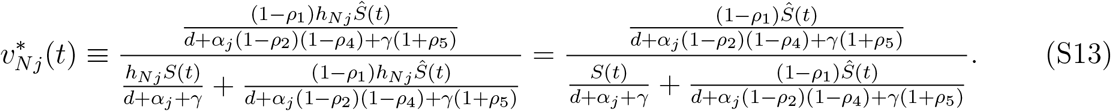

Over long timescales (*t* → ∞), if strain *Nj* is maintained by selection and mutation is small then 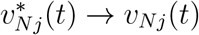; if strain *Nj* is instead maintained by mutation, then although selection (and the approximation (S13)) roughly apply, *v*_*NA*_ will be closer to *v*_*NV*_ than predicted by (S13) (due to mutation). Over short-to-medium timescales, although 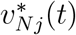 does not exactly match *v*_*Nj*_(*t*) (since this would require *v*_*Nj*_(*t*) to be in quasi-equilibrium), numerical results indicate that the relationship 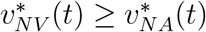 implies *v*_*NV*_ (*t*) ≥ *v*_*NA*_(*t*). It is straightforward to compute

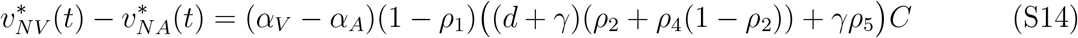

where

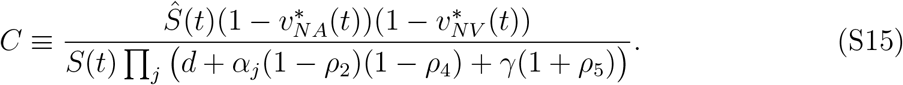

Thus (S14) is positive for all *t*, i.e., 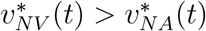, whenever *α*_*V*_ > *α*_*A*_, vaccines imperfectly prevent infection (*ρ*_1_ < 1), and at least one of *ρ*_2_, *ρ*_4_, or *ρ*_5_ are positive. It is easy to see that 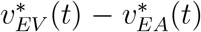 will assume a similar form to (S14), with each of the *ρ*_*i*_ replaced by (1 − *e*)*ρ*_*i*_, *γ* replaced with *γ*(1 + *c*_*γ*_), and 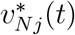 replaced with 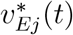; thus we should also expect 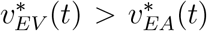 whenever *α*_*V*_ > *α*_*A*_, at least one of *ρ*_2_, *ρ*_4_, or *ρ*_5_ are positive, and *e* < 1 (Fig. S1).

In Figure S1 we provide numerical support that the relation 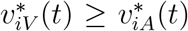 holds over a wide range of parameter space using three different initial conditions; in all conditions we start from a situation in which there is only strain *NA* present and we allow mutation to introduce variation.

1. For the first set of initial conditions (Fig. S1**a**-**c**), we assume that we start from a vaccinated population at the strain *NA* endemic equilibrium, that is, for a given parameter set, we set *f*_*E*_, *f*_*V*_, *D*, *v*_*NV*_, *v*_*EV*_, *v*_*EA*_ to zero in (S4), (S8), (S9) and (S10), and solve for the (positive) values of *S*, 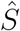, *v*_*NA*_, *I*_*T*_.
2. For the second set of initial conditions (Fig. S1**d**-**f**), we assume the population is at the strain *NA* endemic equilibrium before vaccination, that is,

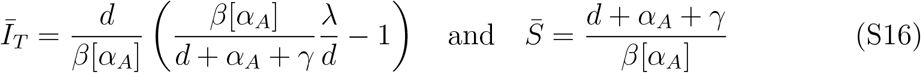

with 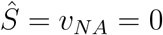.
3. Finally, in the third set of initial conditions (Fig. S1**g**-**i**) we assume at the strain *NA* endemic equilibrium in the unvaccinated population (equation (S16)), we vaccinate a fraction *p* of the uninfected hosts; therefore we have 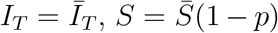 and 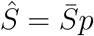, and *v*_*NA*_ = 0, where 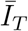 and 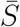 are given in (S16).

For all three initial conditions, we see the prediction that 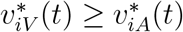 holds (Fig. S1).

### S1.3 Evolution of vaccine-escape

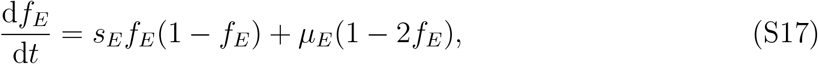

where the selection coefficient for vaccine escape, *s*_*E*_, is

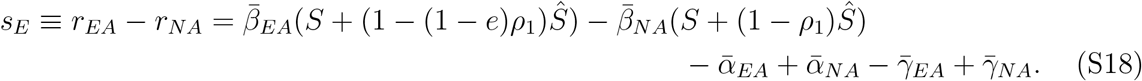

By inspection of *s*_*E*_, decreasing the costs of escape (*c*_*β*_, *c*_*γ*_) increases transmissibility and duration of carriage of strain *EA*, favouring allele *E*. Likewise, increasing the vaccine protection against infection (*ρ*_1_), growth (*ρ*_2_), transmission (*ρ*_3_) and clearance (*ρ*_5_), reduces strain *NA* fitness and so favours allele *E*. Vaccine protection against virulence (*ρ*_4_) is an exception as it increases the duration of carriage without a concomitant reduction in transmission, and so increases strain *NA* fitness relative to strain *EA*. Increasing the degree of escape (*e* → 1) amplifies the selective consequences of the *ρ*_*i*_ by decreasing their effect upon strain *EA* relative to strain *NA*. This can increasingly favour (*ρ*_1_, *ρ*_2_, *ρ*_3_, *ρ*_5_), or disfavour (*ρ*_4_), allele *E*.

### S1.4 Evolution of virulence

If evolution is restricted to the virulence locus (so *μ*_*E*_ = 0 and *f*_*E*_ = *D* = 0), then system (1) reduces to

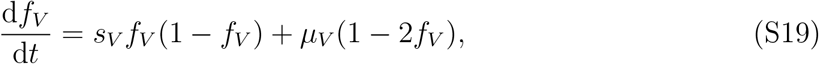

where the selection coefficient for virulence, *s*_*V*_, is

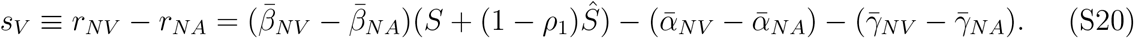

Since the 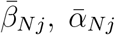 and 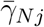 depend upon the *v*_*Nj*_, selection for virulence on the genetic background *N* is a weighted average across two environments: unvaccinated hosts, in which allele *A* is favoured, and vaccinated hosts, in which allele *V* is favoured [11–13]. The weights are the fraction of *Nj*-infections in unvaccinated and vaccinated hosts, that is, 1 − *v*_*Nj*_ and *v*_*Nj*_, respectively (see equation (S6)). Because infections in vaccinated hosts are cleared more rapidly (*ρ*_5_) and may have reduced virulence (*ρ*_2_, *ρ*_4_), higher virulence (and transmission) is favoured in vaccinated hosts. Therefore, allele *A* (lower virulence and transmission) may be favoured in unvaccinated hosts, while allele *V* (higher virulence and transmission) may be favoured in vaccinated hosts [11–13, 15]. Whether allele *A* or *V* is favoured will generally depend upon the vaccine coverage (*p*), the degree to which vaccinated hosts are protected from infection (*ρ*_1_), the transmissibility of infections in vaccinated hosts (*ρ*_3_), as well as the relative difference between *α*_*A*_ and *α*_*V*_ (and *β*[*α*_*A*_] and *β*[*α*_*V*_]; [11–13, 15]).

For example, if we assume that allele *A* is optimal in unvaccinated hosts (or in an entirely unvaccinated population) and allele *V* is optimal in vaccinated hosts (or in an entirely vaccinated population), and take *β*[*α*] = *α^b^*, then

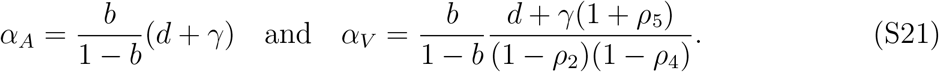

In this circumstance, increasing coverage (*p*) and/or decreasing the vaccines ability to block infection (*ρ*_1_), increases the frequency of infections in vaccinated hosts, *v*_*Nj*_, while decreasing the vaccines ability to block transmission (*ρ*_3_), increases the reproductive value of infections in vaccinated hosts; in either case, this favours allele *V*. On the other hand, increasing the vaccines ability to block growth (*ρ*_2_), to reduce virulence (*ρ*_4_), and/or to clear infections (*ρ*_5_) has two conflicting consequences: on the one hand, it reduces the fitness of *NA*-infections in vaccinated hosts, favouring allele *V*, while on the other hand it increases *α*_*V*_, thereby reducing the fitness of *NV*-infections in unvaccinated hosts, favouring allele *A*. Thus whether increasing *ρ*_2_, *ρ*_4_, or *ρ*_5_ favours allele *V* or *A* as defined in (S21) will depend upon the relative importance of infections in unvaccinated and vaccinated hosts, which is determined by *p*, *ρ*_1_, and *ρ*_3_.

Of course other choices of *α*_*A*_, *α*_*V*_, and *β*[*α*] are possible. However, as we are interested in cases in which *α*_*A*_ is ‘fitter’ in the unvaccinated population on genetic background *N*, and *α*_*V*_ is ‘fitter’ in an entirely vaccinated population on genetic background *N*, for all numerical simulations, we will assume *β*[*α*] = *α*^*b*^ for *b* ∈ (0, 1) and that *α*_*A*_ and *α*_*V*_ are given in (S21).

### S1.5 Epistasis

Epistasis is generated whenever the fitness of an allele depends upon its genetic background, and for continuous time models is defined as

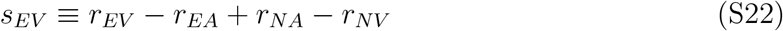

(see e.g., [42, 50]). We are primarily interested in how the parameters controlling vaccine protection contribute to epistasis, and in particular, whether their epistatic contribution is positive or negative. There are three distinct ways in which these contributions can occur: (i) directly, by the parameter of interest explicitly appearing in (S22), (ii) indirectly, by creating or amplifying differences between *α*_*V*_ and *α*_*A*_ (and *β*[*α*_*V*_] and *β*[*α*_*A*_]), and (iii) by affecting the population structure (*v*_*ij*_(*t*)), and/or availability of susceptible hosts (*S*(*t*) and 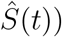. Note that for this section we will explicitly write out the dependence of the variables upon *t* to indicate that epistasis can temporally vary.

In the interests of clarity, we first will assume that vaccine escape is complete (i.e., *e* = 1), before considering how partial vaccine escape affects epistasis. We will assume that *β*[*α*] = *α^b^* which means we can write *β*[*xy*] = *β*[*x*]*β*[*y*]. When *e* = 1, from (S22), the direct contributions (and the sign of the direct contributions) of the vaccine protections are as follows:

#### (1) Vaccines that reduce transmission (*ρ*_1_, *ρ*_2_, *ρ*_3_) produce positive epistasis

Applying the assumption *β*[*α*(1 − *ρ*_2_)] = *β*[*α*]*β*[1 − *ρ*_2_], from (S22) the epistatic contribution of vaccine protection against transmission can be written

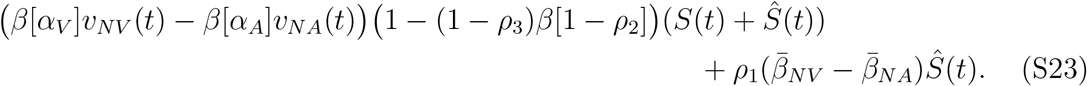

Since *β*[*α*_*V*_] > *β*[*α*_*A*_], whereas we should generally expect that *v*_*NV*_ (*t*) ≥ *v*_*NA*_(*t*) (see Fig. S1) and 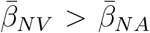 (i.e., strain *NV* is more transmissible then strain *NA*; we should expect this to hold whenever *s*_*V*_ > 0), equation (S23) will be positive, and so vaccines that reduce transmission contribute positive epistasis.

The logic here is that transmission reducing vaccines lower the fitness of allele *N* relative to allele *E*. On the genetic background *N*, this disproportionately affects allele *V* relative to allele *A* because: (i) allele *V* is more transmissible (and the parameters *ρ*_1_, *ρ*_2_, *ρ*_3_ multiplicatively interact with the quantity *β*[*α*]), and (ii) a greater proportion of strain *NV* infections are in vaccinated hosts (*v*_*NV*_ (*t*) ≥ *v*_*NA*_(*t*)). By disproportionately reducing the fitness (equivalently, percapita growth) of strain *NV*, from (S22) this creates positive epistasis.

#### (2) Vaccines that reduce virulence mortality (*ρ*_2_, *ρ*_4_) produce negative epistasis

From (S22), the epistatic contribution of vaccine protection against virulence mortality is

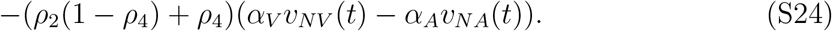

Since *α*_*V*_ > *α*_*A*_ and *v*_*NV*_ (*t*) ≥ *v*_*NA*_(*t*), equation (S24) is negative and so vaccines reducing virulence mortality contributes negative epistasis.

The logic here is that, ignoring any concomitant effects to transmission (by *ρ*_2_), vaccines which reduce virulence mortality increase the fitness of allele *N* relative to allele *E* by increasing infection duration. On the genetic background *N*, this increase in fitness disproportionately affects allele *V* relative to allele *A* because: (i) allele *V* is more virulent (and the parameters *ρ*_2_, *ρ*_4_ multiplicatively interact with *α*_*j*_), and (ii) a greater proportion of strain *NV* infections are in vaccinated hosts (*v*_*NV*_ (*t*) ≥ *v*_*NA*_(*t*)). By disproportionately increasing the fitness (per-capita growth) of strain *NV*, from (S22), this creates positive epistasis.

#### (3) Vaccines that increase clearance (*ρ*_5_) produce positive epistasis

The epistatic contribution of vaccines hastening clearance is

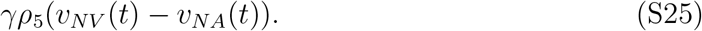

Since *v*_*NV*_ (*t*) ≥ *v*_*NA*_(*t*), equation (S25) is negative, and so vaccines increasing clearance contribute negative epistasis.

The logic here is that vaccines which increase clearance reduce the fitness of allele *N* relative to allele *E* by decreasing infection duration. On the genetic background *N*, this decrease in fitness disproportionately affects allele *V* relative to allele *A* because a greater proportion of strain *NV* infections are in vaccinated hosts (*v*_*NV*_ (*t*) ≥ *v*_*NA*_(*t*)). By disproportionately decreasing the fitness (per-capita growth) of strain *NV*, from (S22), this creates negative epistasis.

In addition to the three categories of epistatic contributions of the vaccine protections, which were also highlighted in the main text, the costs of escape can also produce epistasis. Specifically:

#### (4) Costs of escape to transmission (*c*_*β*_) produces negative epistasis

The epistatic contribution of costs of escape is

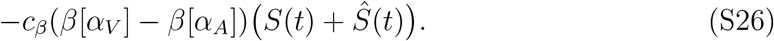

Since *β*[*α*_*V*_] > *β*[*α*_*A*_], equation (S26) is negative, and so costs of escape to transmission contributes negative epistasis.

The logic here is that costs of escape reduce the fitness of allele *E* relative to allele *N* by blocking transmission. On the genetic background *E*, this decrease in fitness disproportionately affects allele *V* relative to allele *A* because allele *V* is more transmissible and the costs of transmission interact multiplicatively with *β*[*α*]. By disproportionately decreasing the fitness (per-capita growth) of strain *EV*, from (S22) this creates negative epistasis.

The costs of escape to duration of carriage, *c*_*γ*_, do not directly contribute epistasis because when vaccine escape is complete, they neither interact directly with the virulence locus (i.e., through multiplicative interactions with *α*_*j*_ and/or *β*[*α*]) nor are they influenced by population structure (i.e., the difference between *v*_*iA*_(*t*) and *v*_*iV*_ (*t*)). Therefore when *e* = 1, the sum of equations (S23), (S24), (S25), and (S26) is equal to (S22), and so the above accounts for all possible direct epistatic contributions.

#### S1.5.1 Influence of partial vaccine-escape on epistasis

In the above, we assumed vaccine-escape was complete, *e* = 1, for clarity. It is of interest to understand how incomplete vaccine-escape affects our interpretation of the epistatic contribution of each vaccine protection. For simplicity, here we consider the epistatic contribution of each of the *ρ*_*i*_ separately (i.e., if we are considering *ρ*_*i*_, then *ρ_j_* = 0, *j* ≠ *i*), and initially assume that the costs of escape are negligible. We also will assume throughout that *β*[*α*] = *α*^*b*^. Define

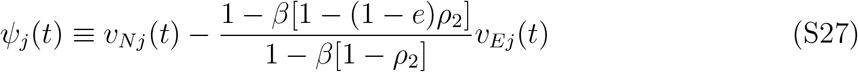

and

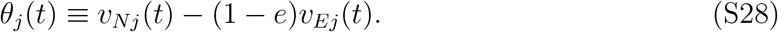

As we will see, allowing for incomplete escape (*e* < 1) yields the same predictions for the sign of the epistatic contribution as when escape is complete (*e* = 1) provided *ψ*_*V*_ (*t*) > *ψ*_*A*_(*t*) and *θ*_*V*_ (*t*) > *θ*_*A*_(*t*). Notice that

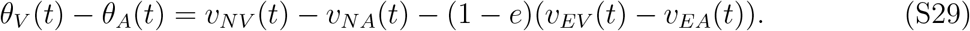

Thus since *v*_*iV*_ (*t*) ≥ *v*_*iA*_(*t*) (see Sup. Info. S1.2.1), and given that for 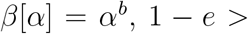 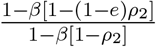 if *θ*_*V*_ (*t*) > *θ*_*A*_(*t*), then *ψ*_*V*_ (*t*) > *ψ*_*A*_(*t*). Further, from (S29), it is clear that as *e* increases, *θ*_*V*_ (*t*) > *θ*_*A*_(*t*) will be satisfied. More generally, if we assume that vaccine escape is complete (*e* = 1), then all else being equal we would expect *v*_*EA*_(*t*) = *v*_*EV*_ (*t*) (it is straightforward to check that 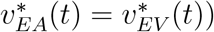. On the other hand, if we assume that as vaccine escape becomes more incomplete (*e* → 0) it also becomes less costly (*c*_*β*_, *c*_*γ*_ → 0), then all else being equal we should expect *v*_*Ej*_(*t*) → *v*_*Nj*_(*t*) as *e* → 0. In combination, we should generally expect that the difference *v*_*iV*_ (*t*) − *v*_*iA*_(*t*) will be larger when *i* = *N* rather than when *i* = *E*, which would imply *θ*_*V*_ (*t*) > *θ*_*A*_(*t*). Indeed, numerical results show that *θ*_*V*_ (*t*) > *θ*_*A*_(*t*) generally holds, even for small *e* (Fig. S2). However, although rare, there are exceptions (Fig. S2); thus all that can be said is that generally speaking *θ*_*V*_ (*t*) > *θ*_*A*_(*t*) (regardless of *e*), and that there is always a level of escape beyond which *θ*_*V*_ (*t*) > *θ*_*A*_(*t*) will hold. In what follows we will assume *θ*_*V*_ (*t*) > *θ*_*A*_(*t*) holds.

With this in mind, the epistatic contribution of each of the *ρ*_*i*_ under incomplete vaccine escape (*e* < 1) are as follows:

1. Vaccines reduce infection, *ρ*_1_ > 0. Here

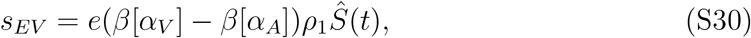

which is **positive** because *β*[*α*_*V*_] > *β*[*α*_*A*_].
2. Vaccines reduce pathogen growth, *ρ*_2_ > 0. In this case,

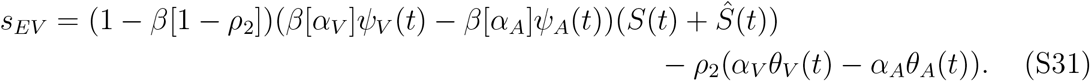

Observe that the effect on transmission (first term) is **positive** since *β*[*α*_*V*_] > *β*[*α*_*A*_] and *ψ*_*V*_ (*t*) > *ψ*_*A*_(*t*), while the effect on virulence (second term) is **negative** since *α_V_ > α_A_* and *θ*_*V*_ (*t*) > *θ*_*A*_(*t*).
3. Vaccines reduce transmission, *ρ*_3_ > 0. Here

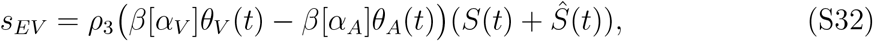

which is **positive** because *β*[*α*_*V*_] > *β*[*α*_*A*_] and *θ*_*V*_ (*t*) > *θ*_*A*_(*t*).
4. Vaccines reduce virulence, *ρ*_4_ > 0. Here

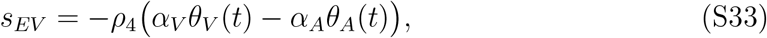

which is **negative** because *α*_*V*_ > *α*_*A*_ and *θ*_*V*_ (*t*) > *θ*_*A*_(*t*).
5. Vaccines increase clearance, *ρ*_5_ > 0. In this case,

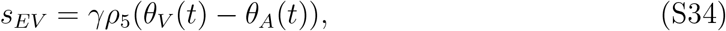

which is **positive** because *θ*_*V*_ (*t*) > *θ*_*A*_(*t*). Thus we see that the inclusion of partial vaccine escape will not change the sign of the epistatic contributions of the different vaccine protections whenever *θ*_*V*_ (*t*) > *θ*_*A*_(*t*); it can, however, reduce the magnitude of the epistatic contribution. In order for epistasis to exist, there must be some difference between genetic backgrounds. Thus the primary influence of the degree of vaccine escape is to alter the magnitude, rather than sign, of the epistatic contributions of each of the vaccine protections by altering the relative difference between genetic backgrounds *E* and *N*. In the event that *θ*_*V*_ (*t*) < *θ*_*A*_(*t*), this will be most likely to change the sign of the epistatic contribution of vaccines affecting clearance, *ρ*_5_; although it can affect *ρ*_2_, *ρ*_3_, *ρ*_4_, this is much less likely because in each of these circumstances *θ_j_*(*t*) is multiplied by either *β*[*α*_*j*_] or *α*_*j*_ and *β*[*α*_*V*_] > *β*[*α*_*A*_] and *α_V_ > α_A_*. Next, how do the costs of vaccine escape contribute to epistasis? Here we allow for all possible vaccine protections to be operating simultaneously.
6. Vaccine-escape reduces transmissibility, *c*_*β*_ > 0. Then the epistatic contribution of the costs of vaccine escape are

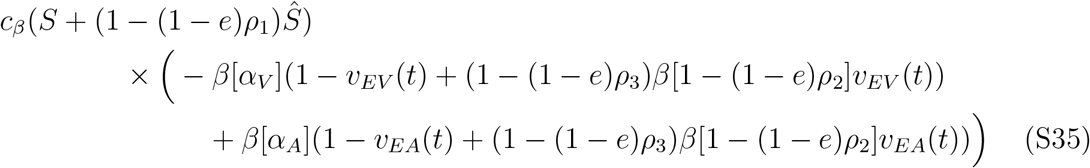 If *e* = 1, this is **negative** (see equation (S26)). If *e* < 1, however, the sign of (S35) is less clear since

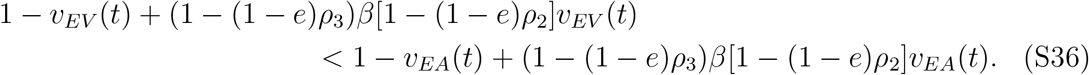 Thus whether the epistatic contribution of *c*_*β*_ is positive or negative may depend in a complex fashion upon the parameter choices, as well as the choice of *β*[*α*].
7. Vaccine-escape reduces duration of carriage, *c*_*γ*_ > 0. Then the epistatic contribution of the costs of vaccine escape are

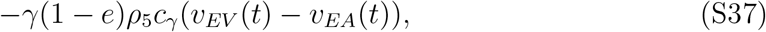

which is **negative** provided *e* < 1 (and zero otherwise).

As we typically lack knowledge about the precise type, or magnitude, of the costs of vaccine escape, and given that how costs are parameterized can lead to different conclusions and evolutionary outcomes in multilocus models [42], in our numerical simulations we assume *e* = 1 as here the costs have a straightforward epistatic contribution.

#### S1.5.2 Influence of virulence *α*_*A*_ and *α*_*V*_ on epistasis

Our analysis assumes that allele *A* is generally ‘fitter’ in unvaccinated hosts (or in an un-vaccinated population) and allele *V* is generally ‘fitter’ in vaccinated hosts (or in an entirely vaccinated population), and so *α*_*V*_ > *α*_*A*_ and *β*[*α*_*V*_] > *β*[*α*_*A*_]. But how does increasing the level of virulence, and transmission, of allele *V* relative to allele *A* affect epistasis? In a similar fashion to partial vaccine escape (*e* < 1), from inspection of (S23), (S24), (S25), and (S26), it is clear that increasing *α*_*V*_ relative to *α*_*A*_ will, all else being equal, increase the magnitude of epistasis, but leave the sign unaffected.

#### S1.5.3 Influence of vaccine coverage on epistasis

Notice that vaccine coverage, *p*, does not directly contribute to epistasis. However, vaccine coverage is very important as it controls the influence of population structure. Specifically, when *p* → 0 or *p* → 1, population structure vanishes. If the population is fully unvaccinated, *p* = 0, then 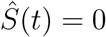 and *v*_*ij*_(*t*) = 0, and so equation (S22) reduces to

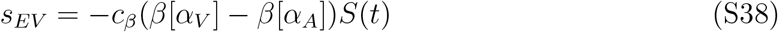

which is negative provided *c*_*β*_ > 0. So in the unvaccinated population, epistasis can only be **negative**.

At the other extreme, when the entire population is vaccinated, *p* = 1, then *S*(*t*) = 0 and *v*_*ij*_(*t*) = 1, and so equation (S22) reduces to

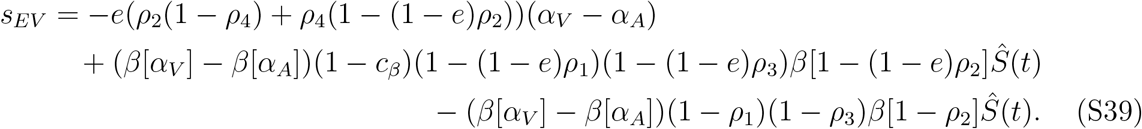

Unlike equation (S38), equation (S39) can be positive or negative, depending upon the protective effects of vaccines relative to the costs of escape. For example, if *ρ*_2_ = *ρ*_4_ = 0, then epistasis will be positive whenever (1 − *c*_*β*_)(1 − (1 − *e*)*ρ*_1_)(1 − (1 − *e*)*ρ*_3_) > (1 − *ρ*_1_)(1 − *ρ*_3_).

Most of the protective effects of vaccination will have the largest impact upon epistasis when population structured is maximised. For example, if vaccine escape is complete (*e* = 1), then from inspection of (S23), (S24), (S25), and (S26), the direct epistatic contributions of the vaccine protections *ρ*_2_, *ρ*_3_, *ρ*_4_, and *ρ*_5_ will be the most substantial as the difference between *v*_*NV*_ (*t*) and *v*_*NA*_(*t*) grows. Since for *p* = 0 or *p* = 1, *v*_*iA*_(*t*) = *v*_*iV*_ (*t*), it follows that the difference between *v*_*NV*_ (*t*) and *v*_*NA*_(*t*) will be greatest for intermediate vaccination rates (intermediate *p*). Thus intermediate vaccination rates will maximise the direct epistatic contributions of the vaccine protections *ρ*_2_, *ρ*_3_, *ρ*_4_, and *ρ*_5_. On the other hand, the (positive) epistatic contribution of *ρ*_1_ will generally be the greatest as vaccination rate increases, as although the impact of *ρ*_1_ depends upon the difference between *v*_*NV*_ (*t*) and *v*_*NA*_(*t*), its effect is also directly proportional to 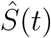, which increases with *p*.

### S1.6 Negative epistasis and evolutionary bistability

Negative epistasis occurs whenever the main vaccine protections are to reduce virulence mortality (*ρ*_2_, *ρ*_4_). These results are generally robust if vaccine protections also include an effect on *ρ*_1_, *ρ*_3_, and *ρ*_5_, provided these effects are smaller than the effects on *ρ*_2_ and *ρ*_4_.

Assume both traits are under directional selection (*s*_*E*_ > 0, *s*_*V*_ > 0) and epistasis is negative. Then negative LD will build up in the population; specifically, the population will tend to be composed of strains *EA* and *NV*. Thus the escape and virulence adaptations are in direct competition with one another. From the strain *NA* equilibrium, initially each adaptation exerts a limited influence upon one another, allowing both alleles *E* and *V* to increase in frequency. However, as *f*_*E*_ + *f*_*V*_ → 1, direct competition between strains *EA* and *NV* sets in and any further increase in allele *E* or *V* comes at the expense of the other. This leads to the population evolving along (or in the vicinity of) the curve

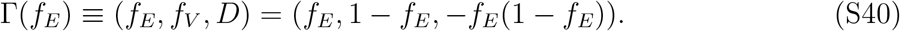

In the vicinity of curve Γ(*f*_*E*_), and assuming mutation and recombination rates are sufficiently small (i.e., *μ*_*E*_, *μ*_*V*_, and *σ* small), system (S8) reduces to

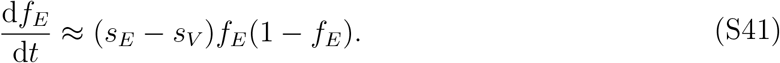

Therefore whichever allele has the larger additive selection coefficient (i.e., *s*_*E*_ or *s*_*V*_), will outcompete the other.

When recombination is frequent, then LD will be continuously removed from the population. Consequently, once the system is in the vicinity of the curve Γ(*f*_*E*_), recombination will ‘push’ the population towards the equilibria along the curve that the population can reach without LD increasing. This can be seen by supposing that we are in the vicinity of (say) the strain *EA* equilibrium, and so *f*_*E*_ ≈ 1 and *f*_*V*_ ≈ 0. While this holds, the dynamical equation for LD can be approximated by

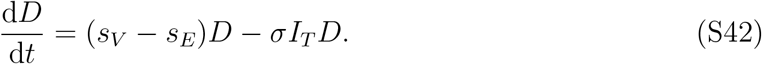

Suppose *s*_*V*_ > *s*_*E*_, that is, selection favours strain *NV* over *EA*, and *D* < 0 (but small). In order for the population to transition from the (*f*_*E*_, *f*_*V*_, *D*) = (1, 0, 0) equilibrium to the (*f*_*E*_, *f*_*V*_, *D*) = (0, 1, 0) equilibrium, significant negative LD must build-up in the population. But from (S42) we see that although selection favours an increase in LD, recombination acts to reduce it. Thus if recombination is sufficiently frequent to overwhelm selection, strain *NV* cannot evolve, despite being selectively fitter than strain *EA*. This bistability induced by recombination creates an advantage for faster growth, as recombination will tend to ‘lock-in’ this advantage in the long-term, when in the absence of recombination, long-term selection would favour an alternate strain.

This bistability is distinct from the one documented in single-locus models [11, 12]. In those models, bistability occurred at the virulence locus because pathogens could either evolve to specialize in unvaccinated hosts (low virulence) or specialize in vaccinated hosts (high virulence); as virulence was modelled as a continuous trait [11, 12], the population evolved towards the more accessible of the two fitness ‘peaks’ from the wild-type virulence. In contrast, here the fitness ‘peaks’ correspond to the different adaptations (strain *EA* vs. strain *NV*), and the fitness ‘valley’ separating the peaks is generated by negative epistasis between alleles *E* and *V*. Recombination breaks up negative LD, preventing the population from crossing the fitness valley, and so creates an evolutionary bistability. The potential for recombination to create an evolutionary bistability has also been observed in models of multidrug resistance [41]. In that case, it prevented the loss of resistance in the absence of treatment, but the principle was similar.

### S1.7 Positive epistasis and the evolution of virulence

Positive epistasis is produced whenever the primary vaccine protections are to reduce infection/transmission (*ρ*_1_, *ρ*_3_) and/or to increase clearance (*ρ*_5_).

Assume both traits are under directional selection (*s*_*E*_ > 0 and *s*_*V*_ > 0) and epistasis is positive. Then there are two scenarios for the transient dynamics: when one selection coefficient is much greater than the other in magnitude, and when the two selection coefficients are of similar strength. Consider the former scenario, and assume that *s*_*E*_ ≫ *s*_*V*_ (*s*_*V*_ ≫ *s*_*E*_ follows similarly). In this case, system (S8) can be approximated by

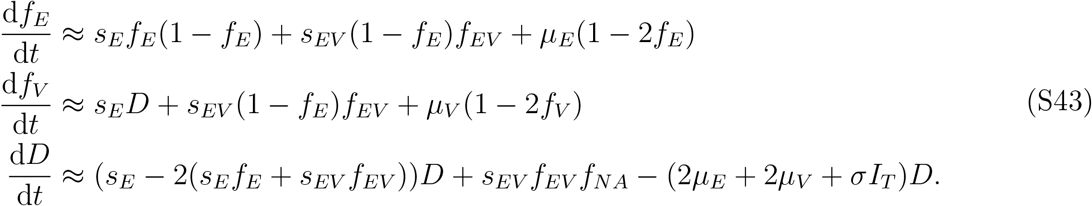

Thus from (S43), the escape allele will rapidly increase in frequency in the population due to directional selection (*s*_*E*_), irrespective of its genetic background (allele *A* or *V*), while the rate of increase in allele *V* (relative to allele *E*) will depend primarily upon the amount of LD present. Specifically, whichever genetic background allele *E* is most commonly found on initially will dominate; thus hitch-hiking occurs at the virulence locus [54]. As a consequence, whichever of strain *EA* or *EV* has the initial numerical advantage will tend to dominant transiently (Fig. S3):

- If the strains *EA*, *EV*, and *NV* are initially absent and are generated by independent mutations at either loci, then we would expect strain *EA* to initially have the numerical advantage over strain *EV* from the strain *NA* equilibrium, as it is accessible by a single mutation. This equally applies if the population is initially in a state at which LD is zero, *D*(0) = 0, in the vicinity of the strain *NA* equilibrium (and so strains *EA*, *EV*, and *NV* are present, but rare); in this circumstance, strain *EA* (although rare) must be initially much more abundant than strain *EV* (to ensure *D*(0) = 0 since *f*_*NA*_ ≈ 1) (Fig. S3**a**).
- If instead strain *EA* and *EV* are initially equally abundant (but rare) or double-mutations (e.g., strain *NA* mutating to strain *EV*) occur at a comparable rate to single mutations, then because epistasis is positive, while (weak) directional selection favours allele *V*, strain *EV* will transiently increase and substantial positive LD will build up in the population (Fig. S3**c**).

Thus if *s*_*E*_ ≫ *s*_*V*_, even though (positive) epistasis produces positive LD, since *s*_*E*_ ≫ *s*_*V*_, whichever of strains *EA* or *EV* has the initial numerical advantage will dominate the transient dynamics. Alternatively, selection favouring the escape allele is of equal strength to selection favouring the virulence allele (i.e. *s*_*E*_ ≈ *s*_*V*_). In this case, both allele *E* and *V* will increase at a similar rate in the population and so owing to the positive epistasis, substantial positive LD can transiently build up (Fig. S3**c**).

Although a variety of factors impact the strength of selection (the relative magnitude of *s*_*E*_ to *s*_*V*_; see Sup. Info. S1.3 and Sup. Info. S1.4), one key mechanism is the costs of escape (*c*_*γ*_, *c*_*β*_) and the degree of escape (*e*). For example, if escape is near complete (*e* → 1), then *c*_*β*_ will produce negative epistasis, while *c*_*γ*_ has no epistatic contribution. So suppose *c*_*β*_ = 0, and consider manipulating *c*_*γ*_. When *c*_*γ*_ is small, escape is virtually cost-free, and so *s*_*E*_ ≫ *s*_*V*_. As *c*_*γ*_ increases, however, escape becomes more costly, until *s*_*E*_ ≈ *s*_*V*_. Finally, increasing *c*_*γ*_ further will eventually lead to *s*_*V*_ ≫ *s*_*E*_.

Next, consider what happens in the long-term. Eventually one (or both) of alleles *E* and *V* will go to quasi-fixation due to directional selection. Suppose allele *E* is more strongly selected for than allele *V*, and so reaches quasi-fixation first; the case when *V* is more strongly selected for follows similarly. Once *f*_*E*_ → 1, LD will necessarily disappear (i.e., *D* → 0), and so the dynamics of system (S8) reduces to

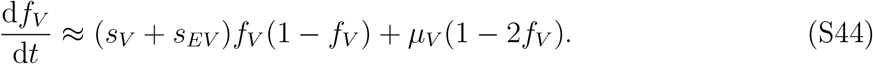

Thus the long-term evolution of virulence on genetic background *E* is determined by the sign of *s*_*V*_ + *s*_*EV*_ = *r*_*EV*_ − *r*_*EA*_. Thus allele *V* (say) can reach high abundances transiently, but ultimately be disfavoured in the long-term.

As an example, consider the case in which vaccine escape is roughly complete, *e* → 1. Then the population structure induced by vaccination does not affect the virulence allele when *f*_*E*_ ≈ 1. Therefore evolution of virulence on the genetic background *E* is a maximisation problem, that is, if we define

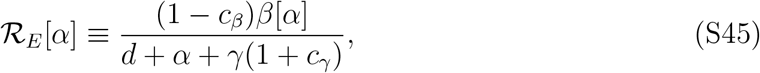

then if 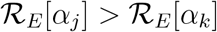, strain *Ej* is competitively superior to strain *Ek*. Solving 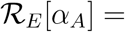 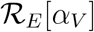 for 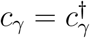 gives

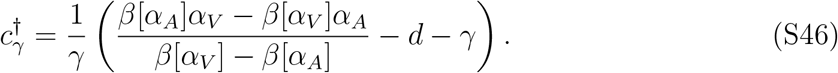

If 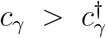, higher virulence is selected for on the genetic background *E*. The logic is straightforward: increasing *c*_*γ*_ increases the clearance rate of *Ej*-infections, selecting for a compensatory increase in virulence.

In fact, higher virulence than the optimal virulence on the genetic background *N* (or the optimal virulence on the genetic background *N* in an entirely vaccinated population, see equation (S21)) can be favoured on the genetic background *E*. Suppose *β*[*α*] = *α*^*b*^, and denote *α*_*E*_ as the *α* maximising 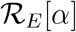, that is,

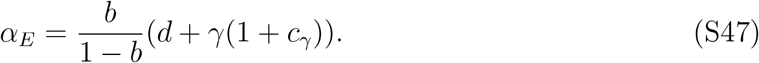

Thus the critical parameter controlling the optimal virulence on the genetic background *E* is the costs of escape to duration of carriage, *c*_*γ*_. If we set *α*_*E*_ = *α*_*V*_ and solve for 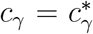, this gives

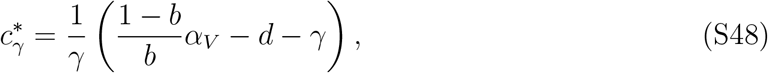

where *α*_*V*_ is given by equation (S21). Hence if 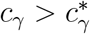, selection favours higher virulence on the genetic background *E* (i.e., *α*_*V*_ < *α*_*E*_). In general, the likelihood that 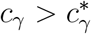 decreases with more effective vaccines (higher *ρ*_2_, *ρ*_4_ and *ρ*_5_) as this will increase *α*_*V*_ (see equation (S21)), reducing the likelihood that *α*_*E*_ > *α*_*V*_. Thus if vaccine escape is sufficiently costly to duration of carriage, a higher level of virulence than that which maximises fitness in vaccinated hosts can be selected for.

### S1.8 Recombination

If recombination rate is large relative to the additive selection coefficients and epistasis, i.e., *σ* ≫ *s*_*E*_, *s*_*V*_, *s*_*EV*_, then the dynamics of LD occurs on a fast timescale relative to change in allele frequencies, and so LD can be assumed to be in a quasi-steady state, say *D* = *D*_QLE_. This quasi-steady state of LD is referred to as quasi-linkage equilibrium (QLE; [51–53]). If we set d*D*/d*t* = 0 and use a perturbative approximation to solve for *D* = *D*_QLE_, to subleading order we obtain

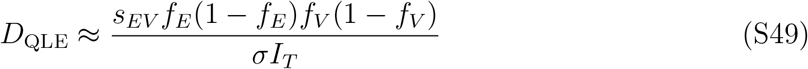

which in turn reduces system (S8) to

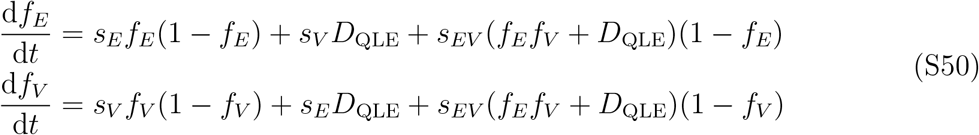

where we have assumed the per-capita mutation rate is very small and so neglected it. If we compute the Jacobian of system (S50) and check the local stability of the *EA* and *NV* - equilibria, the strain *EA* equilibrium will be stable if *s*_*E*_ > 0 and *s*_*V*_ + *s*_*EV*_ < 0, while the strain *NV* equilibrium will be stable if *s*_*V*_ > 0 and *s*_*E*_ + *s*_*EV*_ < 0. Given directional selection favouring escape (*s*_*E*_ > 0) and virulence (*s*_*V*_ > 0), it is apparent that a bistability will occur if epistasis is negative and strong (i.e., −*s*_*EV*_ > max(*s*_*E*_, *s*_*V*_)). The logic here is that recombination prevents double mutants from rising to abundance, and thus in the vicinity of their respective equilibria, the *EA* and *NV* infections are only competing against the *EV* and *NA* strains and not each other. Since epistasis is negative and directional selection favours alleles *V* and *E*, strain *EV* and strain *NA* will tend to be a selective disadvantage relative to either strain *EA* or *NV*.

---

**Figure S1:**
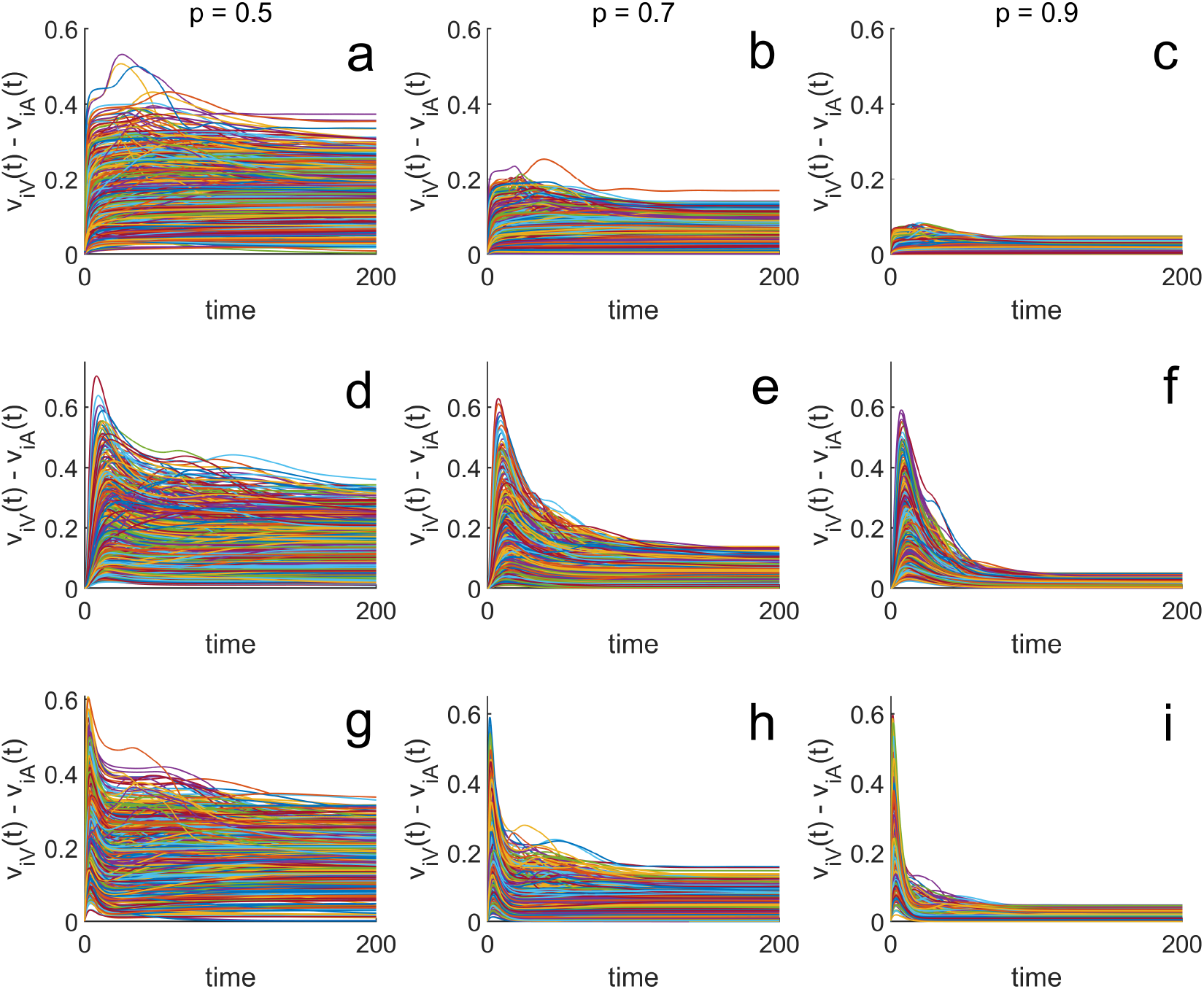
A greater fraction of *iV* infections are in vaccinated hosts than *iA* infections, *v*_*iV*_ (*t*) ≥ *v*_*iA*_(*t*). Each panel shows the difference between *v*_*iV*_ (*t*) and *v*_*iA*_(*t*) for 250 randomly chosen parameter sets with the fraction vaccinated (*p*) given above each column. Note that in all cases, as expected (see Sup. Info. S1.2.1), *v*_*iV*_ (*t*) ≥ *v*_*iA*_(*t*). Each row corresponds to a different set of initial conditions (ICs) described in Sup. Info. S1.2.1: the first row corresponds to ICs (1), the second to ICs (2), and the third to ICs (3). For each simulation, 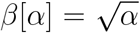, with *α*_*A*_ and *α*_*V*_ as in equation (S21), while *λ* = *d* = 0.05, *γ* = 0.1, *μ*_*E*_ = 10^−4^, *μ*_*V*_ = 10^−4^, *σ* = 0, and the remaining parameters are chosen uniformly at random over the given intervals: (1 − *e*), *ρ*_1_, *ρ*_2_, *ρ*_3_, *ρ*_4_, *c*_*β*_ ∈ [0, 0.95], *ρ*_5_ ∈ [0, 8], *c*_*γ*_ ∈ [0, 2]. Hence, each simulation start with a monomorphic pathogen population (only the *NA* genotype is present initially) and mutation introduces genetic variation and allows pathogen adaptation to vaccination. All parameter sets satisfied the constraints that *s*_*E*_ > 0 and *s*_*V*_ > 0 at the strain *NA* endemic equilibrium in a vaccinated population (ICs (1), see Sup. Info. S1.2.1), and that *α*_*V*_/*α*_*A*_ > 1.2.

**Figure S2:**
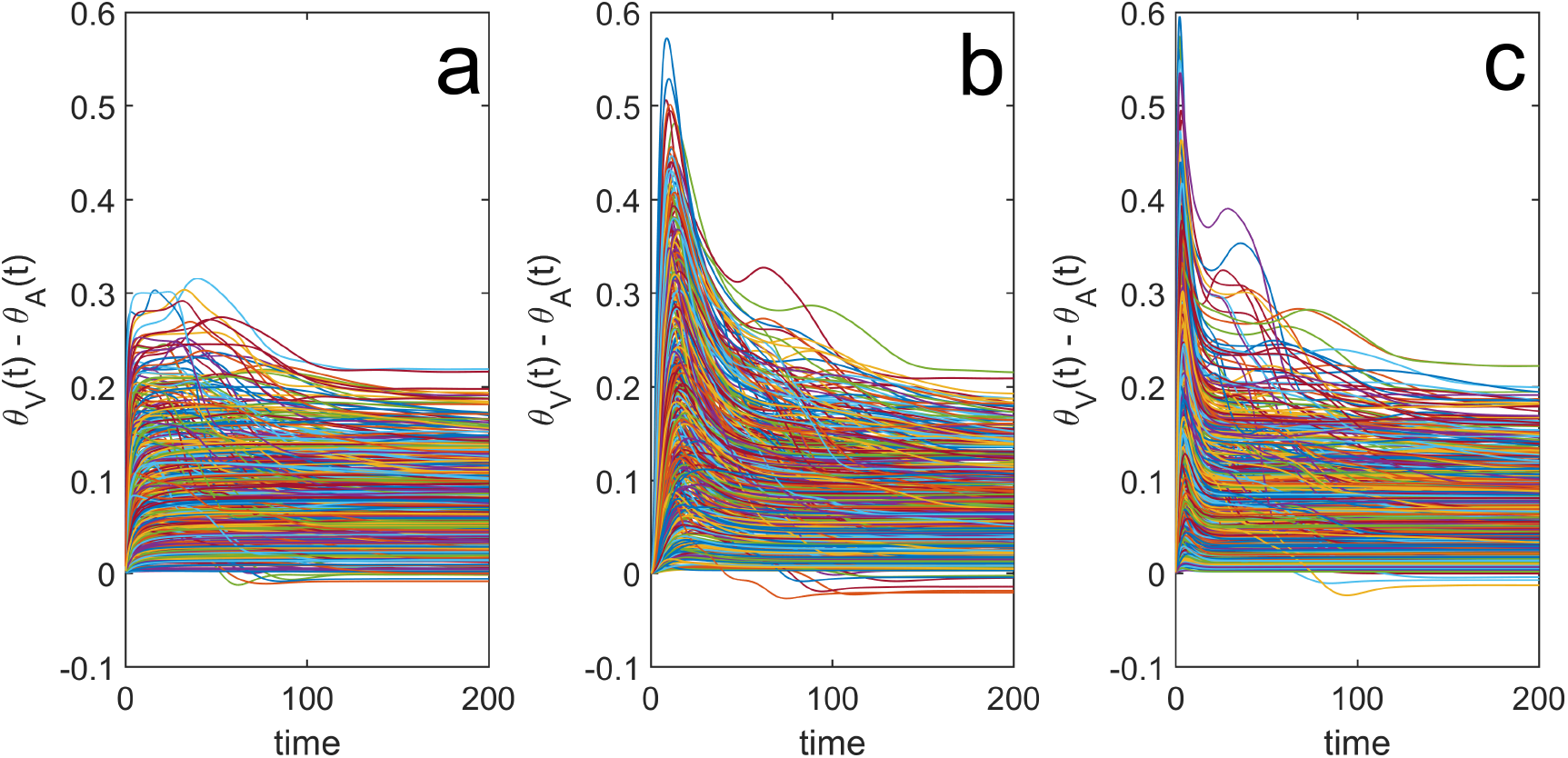
Relationship between *θ*_*V*_ (*t*) and *θ*_*A*_(*t*) when vaccine-escape is incomplete. For almost all simulations, *θ*_*V*_ (*t*) ≥ *θ*_*A*_(*t*); the few simulations that do not satisfy this condition are associated with limited vaccine-escape (small *e*; see Sup. Info. S1.5). Each panel shows the simulation results of 500 randomly chosen parameter sets for a different set of initial conditions (ICs) described in Sup. Info. S1.2.1: panel **a** corresponds to ICs (1); panel **b** to ICs (2); and panel **c** to ICs (3). For all simulations, 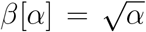, *α*_*A*_ and *α*_*V*_ are given by equation (S21), with *p* = 0.6, *λ* = *d* = 0.05, *γ* = 0.1, *μ*_*E*_ = 10^−4^, *μ*_*V*_ = 10^−4^, *σ* = 0, and the remaining parameters chosen uniformly at random over the intervals: (1 − *e*), *ρ*_1_, *ρ*_2_, *ρ*_3_, *ρ*_4_, *c*_*β*_ ∈ [0, 0.95], *ρ*_5_ ∈ [0, 8], *c*_*γ*_ ∈ [0, 2]. Finally, all parameter sets satisfied the constraints that *s*_*E*_ > 0 and *s*_*V*_ > 0 at the strain *NA* endemic equilibrium in a vaccinated population (ICs (1), see Sup. Info. S1.2.1), and that *α*_*V*_/*α*_*A*_ > 1.2.

**Figure S3:**
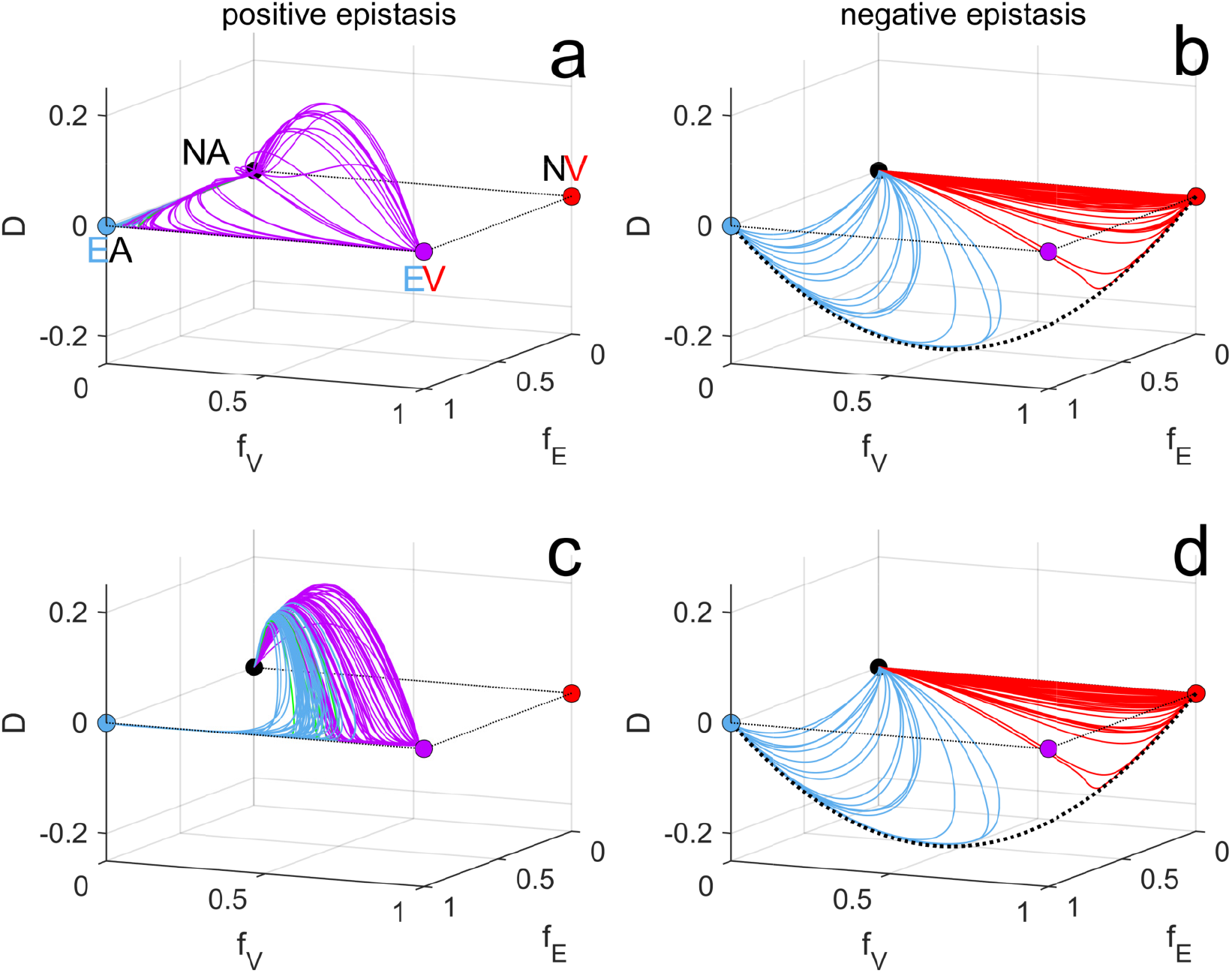
When epistasis is positive, the evolutionary dynamics are sensitive to initial conditions. Panels **a**, **c** are when epistasis is positive, while panels **b**, **d** are when epistasis is negative. In panels **a**, **b**, double-mutations are not possible (e.g., strain *NA* cannot mutate to strain *EV* and vice-versa), whereas in panels **a**, **d** double mutations occur at the same per-capita rate as single mutations. If epistasis is positive (panel **d**), this leads to a substantial, transient build-up of LD, whereas if epistasis is negative (panel **c**), this has negligible effect. Each panel shows 100 simulations starting from a monomorphic pathogen population (only the *NA* genotype is present initially) at it’s endemic equilibrium following vaccination; mutation introduces genetic variation and allows pathogen adaptation to vaccination. Parameters used were 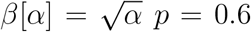, *e* = 1, *λ* = *d* = 0.05, *γ* = 0.1, *μ*_*E*_ = 10^−4^, *μ*_*V*_ = 10^−4^, *σ* = 0. For panels **a**, **c**, *ρ*_2_ = *ρ*_4_ = *c*_*β*_ = 0, while *ρ*_1_, *ρ*_3_, *ρ*_5_, *c*_*γ*_ were chosen uniformly at random from the intervals [0, 0.95], [0, 0.95], [0, 15], [0, 5], respectively. For panels **b**, **d**, *ρ*_1_ = *ρ*_3_ = *ρ*_5_ = 0, while *ρ*_2_, *ρ*_4_, *c*_*β*_, *c*_*γ*_ were chosen uniformly at random from the intervals [0, 0.95], [0, 0.95], [0, 0.1], [0, 0.2], respectively. All parameter sets satisfied the constraints that *s*_*E*_ > 0 and *s*_*V*_ > 0 at the strain *NA* equilibrium and *α*_*V*_/*α*_*A*_ > 1.25. *α*_*A*_ and *α*_*V*_ were chosen to be the optimal virulence in an entirely unvaccinated and vaccinated population, respectively (Sup. Info. S1.4).

## Notes

### Competing Interest Statement

The authors have declared no competing interest.

## References

[1] McLean, A. R., 1998 Vaccines and their impact on the control of disease. British Medical Bulletin 54, 545–556.

[2] Kennedy, D. A. & Read, A. F., 2017 Why does drug resistance readily evolve but vaccine resistance does not? Proc. R. Soc. B 284.

[3] Kennedy, D. A. & Read, A. F., 2018 Why the evolution of vaccine resistance is less of a concern than the evolution of drug resistance. Proc. Natl. Acad. Sci. 115, 12878–12886.

[4] Klausner, R. D., Fauci, A. S., Corey, L., Nabel, G. J., Gayle, H., Berkley, S., Haynes, B. F., Baltimore, D., Collins, C., Douglas, R. G. et al., 2003 Enhanced: The need for a global HIV vaccine enterprise. Science 300, 2036–2039.

[5] Morris, D. H., Gostic, K. M., Pompei, S., Bedford, T., Luksza, M., Neher, R. A., Grenfell, B. T., Lassig, M. & McCauley, J. W., 2018 Predictive modeling of influenza shows the promise of applied evolutionary biology. Trends Microbiol. 26, 102–118.

[6] Luksza, M. & Lassig, M., 2014 A predictive fitness model for influenza. Nature 507, 57–61.

[7] Barrat-Charlaix, P., Huddleston, J., Bedford, T. & Neher, R. A., 2021 Limited pre-dictability of amino acid substitutions in seasonal influenza viruses. Mol. Biol. Evol. (doi:10.1093/molbev/msab065).

[8] Arinaminpathy, N., Ratmann, O., Koelle, K., Epstein, S. L., Price, G. E., Viboud, C., Miller, M. A. & Grenfell, B. T., 2012 Impact of cross-protective vaccines on epidemio-logical and evolutionary dynamics of influenza. Proc. Natl. Acad. Sci. 109, 3173–3177.

[9] Subramanian, R., Graham, A. L., Grenfell, B. T. & Arinaminpathy, N., 2016 Universal or specific? A modeling-based comparison of broad-spectrum influenza vaccines against conventional, strain-matched vaccines. PLoS Comp. Biol. 12, e1005204.

[10] Viboud, C., Gostic, K., Nelson, M. I., Price, G. E., Perofsky, A., Sun, K., S., T. N., Cowling, B. J., Epstein, S. L. & Spiro, D. J., 2020 Beyond clinical trials: evolutionary and epidemiological considerations for development of a universal influenza vaccine. PLoS Pathogens 16, e1008583.

[11] Gandon, S., Mackinnon, M. J., Nee, S. & Read, A. F., 2001 Imperfect vaccines and the evolution of pathogen virulence. Nature 414, 751–756.

[12] Gandon, S., Mackinnon, M., Nee, S. & Read, A. F., 2003 Imperfect vaccination: some epidemiological and evolutionary consequences. Proc. R. Soc. B. 270, 1129–1136.

[13] Andre, J. & Gandon, S., 2006 Vaccination, within-host dynamics, and virulence evolution. Evolution 60, 13–23.

[14] Atkins, K. E., Read, A. F., Savill, N. J., Renz, K. G., Fakhrul Islam, A. F. M., Walkden-Brown, S. W. & Woolhouse, M. E. J., 2012 Vaccination and reduced cohort duration can drive virulence evolution: Marek’s disease virus and industrialized agriculture. Evolution 67, 851–860.

[15] Ganusov, V. V. & Antia, R., 2006 Imperfect vaccines and the evolution of pathogens causing acute infections in vertebrates. Evolution 60, 957–969.

[16] Williams, P. D. & Day, T., 2008 Epidemiological and evolutionary consequences of targeted vaccination. Mol. Ecol. 17, 485–499.

[17] Bernhauerová, V., 2016 Vaccine-driven evolution of parasite virulence and immune evasion in age-structured population: the case of pertussis. Theor. Ecol. 9, 431–442.

[18] Anderson, R. M. & May, R. M., 1982 Coevolution of hosts and parasites. Parasitology 85, 411–426.

[19] Frank, S. A., 1996 Models of parasite virulence. Quarterly Rev. Biol. 71, 37–78.

[20] Alizon, S., Hurford, A., Mideo, N. & van Baalen, M., 2009 Virulence evolution and the trade-off hypothesis: history, current state of affairs and the future. J. Evol. Biol. 22, 245–259.

[21] Cressler, C. E., McLeod, D. V., Rozins, C., van den Hoogen, J. & Day, T., 2016 The adaptive evolution of virulence: a review of theoretical predictions and empirical tests. Parasitology 143, 915–930.

[22] Witter, R. L., 1997 Increased virulence of Marek’s disease virus field isolates. Avian Diseases 41, 149–163.

[23] Read, A. F., Baigent, S. J., Powers, C., Kgosana, L. B., Blackwell, L., Smith, L. P., Kennedy, D. A., Walkden-Brown, S. W. & Nair, V. K., 2015 Imperfect vaccination can enhance the transmission of highly virulent pathogens. PLoS Biol. 13, e1002198.

[24] Read, A. F. & Mackinnon, M. J., 2008 Pathogen evolution in a vaccinated world. In Evolution in health and disease, pp. 139–52. Oxford University Press.

[25] Mackinnon, M. J. & Read, A. F., 2004 Immunity promotes virulence evolution in a malaria model. PLoS Biol. 2, 1286–1292.

[26] Barclay, V. C., Sim, D., Chan, B. H. K., Nell, L. A., Rabaa, M. A., Bell, A. S., Anders, R. F. & Read, A. F., 2012 The evoutionary consequences of blood-stage vaccination on the rodent malaria *Plasmodium chabaudi*. PLoS Biol. 10, e1001368.

[27] Mooi, F. R., van Loo, I. H. M., van Gent, M., He, Q., Bart, M. J., Heuvelman, K. J., de Greeff, S. C., Diavatopoulos, D., Teunis, P., Nagelkerke, N. et al., 2009 *Bordetella pertussis* strains with increased toxin production associated with pertussis resurgence. Emerg. Infect. Diseases 15, 1206–1213.

[28] McLean, A. R., 1995 Vaccination, evolution and changes in the efficacy of vaccines: a theoretical framework. Proc. Roy. Soc. B 261, 389–393.

[29] Gupta, S., Ferguson, N. M. & Anderson, R. M., 1997 Vaccination and the population structure of antigenically diverse pathogens that exchange genetic material. Proc. R. Soc. B 264, 1435–1443.

[30] Lipsitch, M., 1997 Vaccination against colonizing bacteria with multiple serotypes. Proc. Natl. Acad. Sci. 94, 6571–6576.

[31] Restif, O. & Grenfell, B. T., 2007 Vaccination and the dynamics of immune evasion. J. Roy. Soc. Interface 4, 143–153.

[32] van Boven, M., Mooi, F. R., Schellekens, J. F. P., de Melker, H. E. & Kretzschmar, M., 2005 Pathogen adaptation under imperfect vaccination: implications for pertussis. Proc. Roy. Soc. B. 272, 1617–1624.

[33] Grenfell, B. T., Pybus, O. G., Gog, J. R., Wood, J. L. N., Daly, J. M., Mumford, J. A. & Holmes, E. C., 2004 Unifying the epidemiological and evolutionary dynamics of pathogens. Science 303, 327–323.

[34] Saad-Roy, C. M., Morris, S. E., Metcalf, C. J. E., Mina, M. J., Baker, R. E., Farrar, J., Holmes, E. C., Pybus, O. G., Graham, A. L., Levin, S. A. et al., 2021 Epidemiological and evolutionary considerations of sars-cov-2 vaccine dosing regimes. Science 372, 363–370. (doi:10.1126/science.abg8663).

[35] Cobey, S., Larremore, D. B., Grad, Y. H. & Lipsitch, M., 2021 Concerns about SARS-CoV-2 evolution should not hold back efforts to expand vaccination. Nat. Rev. Immunol. (doi:https://doi.org/10.1038/s41577-021-00544-9).

[36] Kouyos, R. D., Metcalf, C. J. E., Birger, R., Klein, E. Y., Abel zur Wiesch, P., Ankomah, P., Arinaminpathy, N., Bogich, T. L., Bonhoeffer, S., Brower, C. et al., 2014 The path of least resistance: aggressive or moderate treatment? Proc. Roy. Soc. B 281, 20140566. (doi:10.1098/rspb.2014.0566).

[37] Acevedo, M. A., Dillemuth, F. P., Flick, A. J., Faldyn, M. J. & Elderd, B. D., 2019 Virulence-driven trade-offs in disease transmission: a meta-analysis. Evolution 73, 636–647.

[38] Telenti, A., Arvin, A., Corey, L., Corti, D., Diamond, M. S., Garcia-Sastre, A., Garry, R. F., Holmes, E. C., Pang, P. S. & Virgin, H. W., 2021 After the pandemic: perspectives on the future trajectory of COVID-19. Nature 596, 495–504.

[39] Slatkin, M., 2008 Linkage disequilibrium – understanding the evolutionary past and mapping the medical future. Nat. Rev. Genetics 9, 477–485.

[40] Rice, S. H., 2004 Evolutionary Theory: Mathematical and Conceptual Foundations. Sunderland, MA, USA: Sinauer Associates.

[41] Day, T. & Gandon, S., 2012 The evolutionary epidemiology of multilocus drug resistance. Evolution 66, 1582–1597.

[42] McLeod, D. V. & Gandon, S., 2021 Understanding the evolution of multiple drug resistance in structured populations. eLife 10, e65645.

[43] de Visser, J. A. G. M., Cooper, T. F. & Elena, S. F., 2011 The causes of epistasis. Proc. R. Soc. B 278, 3671–3624.

[44] Felsenstein, J., 1965 The effect of linkage on directional selection. Genetics 52, 349–363.

[45] Lewontin, R. C. & Kojima, K., 1960 The evolutionary dynamics of complex polymorphisms. Evolution 14, 458–472.

[46] de Visser, J. A. G. M. & Elena, S. F., 2007 The evolution of sex: empirical insights into the roles of epistasis and drift. Nat. Rev. Genetics 8, 139–149.

[47] Otto, S. P. & Barton, N. H., 1997 The evolution of recombination: removing the limits to natural selection. Genetics 147, 879–906.

[48] Fenner, F. J., 1983 The Florey Lecture, 1983 - Biological control, as exemplified by smallpox eradication and myxomatosis. Proc. R. Soc. B. 218, 259–285.

[49] Muñoz-Alía, M. Á., Nace, R. A., Zhang, L. & Russell, S. J., 2021 Serotypic evolution of measles virus is constrained by multiple co-dominant b cell epitopes on its surface glycoproteins. Cell Reports Medicine 2, 100225.

[50] Kouyos, R. D., Fouchet, D. & Bonhoeffer, S., 2009 Recombination and drug resistance in HIV: population dynamics and stochasticity. Epidemics 1, 58–69.

[51] Neher, R. A. & Shraiman, B. I., 2011 Statistical genetics and evolution of quantitative traits. Rev. Mod. Phys. 83, 1283–1300.

[52] Kimura, M., 1965 Attainment of quasi-linkage equilibrium when gene frequencies are changing by natural selection. Genetics 52, 875–890.

[53] Otto, S. P. & Day, T., 2007 A Biologist’s Guide to Mathematical Modeling in Ecology and Evolution. Princeton University Press.

[54] Maynard Smith, J. & Haigh, J., 1974 The hitch-hiking effect of a favourable gene. Genet. Res. Camb. 23, 23–35.

[55] Lenski, R. E. & May, R. M., 1994 Evolution of virulence in parasites and pathogens: reconciliation between two competing hypotheses. J. Theor. Biol. 169, 253–265.

[56] Day, T. & Proulx, S. R., 2004 A general theory for the evolution of virulence. Am. Nat. 163, E40–E63.

[57] Bolker, B. M., Nanda, A. & Shah, D., 2010 Transient virulence of emerging pathogens. J. Roy. Sci. Interface 7, 811–822.

[58] Harrison, T. J., Hopes, E. A., J., O. C., Zanetti, A. R. & Zuckerman, A. J., 1991 Independent emergence of a vaccine-induced escape mutant of hepatitis B virus. J. Hepatology 13, S105–S107.

[59] Mooi, F. R., van Oirschot, H., Heuvelman, K., van der Heide, H. G. J., Gaastra, W. & Willems, R. J. L., 1998 Polymorphism in the *Bordetella pertussis* virulence factors P.69/Pertactin and pertussis toxin in the Netherlands: temporal trends and evidence for vaccine-driven evolution. Infect. Immun. 66, 670–675.

[60] Gandon, S. & Day, T., 2008 Evidences of parasite evolution after vaccination. Vaccine 26, C4–C7.

[61] Regev-Yochay, G., Amit, S., Bergwerk, M., Lipsitch, M., Leshem, E., Kahn, R., Lustig, Y., Cohen, C., Doolman, R., Ziv, A. et al., 2021 Decreased infectivity following BNT162b2 vaccination: a prospective cohort study in Israel. The Lancet Regional Health - Europe 7, 100150.

[62] Ke, R., Martinez, P. P., Smith, R. L., Gibson, L. L., Achenbach, C. J., McFall, S., Qi, C., Jacob, J., Dembele, E., Bundy, C. et al., 2021 Longitudinal analysis of SARS-CoV-2 vaccine breakthrough infections reveal limited infectious virus shedding and restricted tissue distribution. medRxiv (doi:10.1101/2021.08.30.21262701).

[63] Gilbert, S. C., 2011 T-cell-inducing vaccines - what’s the future. Immunology 135, 19–26.

[64] Schaffer, A. C. & Lee, J. C., 2008 Vaccination and passive immunisation against *Staphy-lococcus aureus*. International Journal of Antimicrobial Agents 32, S71–S78.

[65] Pica, N. & Palese, P., 2013 Toward a universal influenza virus vaccine: prospects and challenges. Annu. Rev. Med. 64, 189–202.

[66] Pardi, N., Hogan, M. J., Porter, F. W. & Weissman, D., 2018 mRNA vaccines – a new era in vaccinology. Nat. Rev. Drug Discovery 17, 261–279.

[67] Saad-Roy, C. M., McDermott, A. B. & Grenfell, B. T., 2019 Dynamic perspectives on the search for a universal influenza vaccine. J. Infect. Dis. 219, S46–S56.

[68] Arinaminpathy, N., Riley, S., Barclay, W. S., Saad-Roy, C. & Grenfell, B., 2020 Population implications of the deployment of novel universal vaccines against epidemic and pandemic influenza. J. R. Soc. Interface 17, 20190879.

[69] Ekiert, D. C., Bhabha, G., Elsliger, M., Friesen, R. H. E., Jongeneelen, M., Throsby, M., Goudsmit, J. & Wilson, I. A., 2009 Antibody recognition of a highly conserved influenza virus epitope. Science 324, 246–251.

[70] Epstein, S. L. & Price, G. E., 2010 Cross-protective immunity to influenza A viruses. Expert Rev. Vaccines 9, 1325–1341.

[71] Yurkovetskiy, L., Wang, X., Pascal, K. E., Tomkins-Tinch, C., Nyalile, T. P., Wang, Y., Baum, A., Diehl, W. E., Dauphin, A., Carbone, C. et al., 2020 Structural and functional analysis of the D614G sars-cov-2 spike protein variant. Cell 183, 739–751.e8. ISSN 0092-8674.

[72] Ozono, S., Zhang, Y., Ode, H., Sano, K., Tan, T. S., Imai, K., Miyoshi, K., Kishigami, S., Ueno, T., Iwatani, Y. et al., 2021 SARS-CoV-2 D614G spike mutation increases entry efficiency with enhanced ACE2-binding affinity. Nat. Comm. 12.

[73] Jangra, S., Ye, C., Rathnasinghe, R., Stadlbauer, D., Personalized Virology Initiative study group, Krammer, F., Simon, V., Martinez-Sobrido, L., Garcia-Sastre, A. & Schotsaert, M., 2021 SARS-CoV-2 spike E484K mutation reduces antibody neutralisation. Lancet Microbe.

[74] Greaney, A. J., Starr, T. N., Barnes, C. O., Weisblum, Y., Schmidt, F., Caskey, M., Gaebler, C., Cho, A., Agudelo, M., Finkin, S. et al., 2021 Mutational escape from the polyclonal antibody response to SARS-CoV-2 infection is largely shaped by a single class of antibodies. bioRxiv (doi:10.1101/2021.03.17.435863).

[75] Peck, K. M., Burch, C. L., Heise, M. T. & Baric, R. S., 2015 Coronavirus host range expansion and Middle East respiratory syndrome coronavirus emergence: biochemical mechanisms and evolutionary perspectives. Annu. Rev. Virol. 2, 95–117.

[76] Jackson, B., Boni, M. F., Bull, M. J., Colleran, A., Colquhoun, R. M., Darby, A. C., Haldenby, S., Hill, V., Lucaci, A., McCrone, J. T. et al., 2021 Generation and transmission of inter-lineage recombinants in the SARS-CoV-2 pandemic. Cell (doi: https://doi.org/10.1016/j.cell.2021.08.014).

[77] Turkahia, Y., Thornlow, B., Hinrichs, A., McBroome, J., Ayala, N., Ye, C., de Maio, N., Haussler, D., Lanfear, R. & Corbett-Detig, R., 2021 Pandemic-scale phylogenomics reveals elevated recombination rates in the SARS-CoV-2 spike region. bioRxiv (doi: 10.1101/2021.08.04.455157).

[78] VanInsberghe, D., Neish, A. S., Lowen, A. C. & Koelle, K., 2021 Recombinant sars-cov-2 genomes circulated at low levels over the first year of the pandemic. Virus Evolution (doi:10.1093/ve/veab059).

[79] UK SAGE, 2021. Can we predict the limits of SARS-CoV-2 variants and their phenotypic consequences? (doi:https://assets.publishing.service.gov.uk/government/uploads/system/uploads/attachmentdata/file/1007566/S1335LongtermevolutionofSARS-CoV-2.pdf).

[80] Polack, F. P., Thomas, S. J., Kitchin, N., Absalon, J., Gurtman, A., Lockhart, S., Perez, J. L., Pérez Marc, G., Moreira, E. D., Zerbini, C. et al., 2020 Safety and efficacy of the BNT162b2 mRNA Covid-19 vaccine. New England Journal of Medicine 383, 2603–2615.

[81] Dagan, N., Barda, N., Kepten, E., Miron, O., Perchik, S., Katz, M. A., Hernan, M. A., Lipsitch, M., Reis, B. & Balicer, R. D., 2021 BNT162b2 mRNA covid-19 vaccine in a nationwide mass vaccination setting. N. Engl. J. Med. 384, 1412–1423.

[82] Thompson, M. G., Burgess, J. L., Naleway, A. L., Tyner, H., Yoon, S. K., Meece, J., Olsho, L. E., Caban-Martinez, A. J., Fowlkes, A. L., Lutrick, K. et al., 2021 Prevention and attenuation of Covid-19 with the BNT162b2 and mRNA-1273 caccines. New England Journal of Medicine 385, 320–329. (doi:10.1056/NEJMoa2107058).

[83] Baker, R. E., Park, S. W., Yang, W., Vecchi, G. A., Metcalf, C. J. E. & Grenfell, B. T., 2020 The impact of COVID-19 nonpharmaceutical interventions on the future dynamics of endemic infections. Proc. Natl. Acad. Sci. 117, 30547–30553.

[84] Antia, R., Levin, B. R. & May, R. M., 1994 Within-host population dynamics and the evolution and maintenance of microparasite virulence. Am. Nat. 144, 457–472.

[85] Ben-Shachar, R. & Koelle, K., 2018 Transmission-clearance trade-offs indicate that dengue virulence evolution depends on epidemiological context. Nat. Comm. 9, 1–11.

